# Mating display plasticity predicts the biogeography of complex communication and population persistence in changing climates

**DOI:** 10.64898/2026.07.21.739722

**Authors:** Noah T. Leith, Jake P. Woods, Kasey D. Fowler-Finn

**Affiliations:** Department of Biology, Saint Louis University; St. Louis, MO, USA; Department of Biological Sciences, University of Pittsburgh; Pittsburgh, PA, USA; Department of Environmental Science, Policy, and Management, University of California, Berkeley; Berkeley, CA, USA; Department of Integrative Biology, University of South Florida; Tampa, FL, USA

**Keywords:** sexual selection, mate choice, phenotypic plasticity, sensory ecology, climate change

## Abstract

Animals often produce complex combinations of signals to enhance display detection, interpretation, and the information conveyed during communication. Combined signals can serve many functions in communication systems, but we know little about how signal combinations initially evolve and why they often vary in complexity across broad geographic gradients. Here, we show that environmentally driven changes in the coordinated production of multiple signals determine whether the signals can interact in functional ways, and thereby shape the capacity for selection to favor increased display complexity. Our behavioral experiment in a focal species of *Schizocosa* wolf spider reveal that shifts to hotter and wetter environments can enable males with exaggerated morphological ornaments to also produce exaggerated courtship behaviors, allowing these signals to interactively affect mating success. Congruently, phylogenetic analyses show that species in hotter and wetter regions of North America have repeatedly evolved more complex courtship displays alongside exaggerated morphological ornaments. Biogeographic analyses further suggest that geographic gradients in display complexity may arise not only from *in situ* evolution, but also from more frequent establishment of species with complex displays in hotter and wetter regions. Finally, we found that species with complex displays were more likely to persist in areas that have faced more severe climate warming and intensified precipitation in the last 50 years—conditions that should enhance the functioning of complex mating displays for reproductive success. Altogether, our findings reveal a novel association between plasticity in inter-signal interactions and variation in the complexity of communication systems across scales of biological organization.

## Main

The evolution of communication systems underlies much of the world’s biodiversity^1–3^. One key adaptation precipitating this diversity is the evolution of complex, muti-component displays^4^. Coordinating the production of multiple signals can be adaptive in communication by allowing the signals to interact in functional ways^5,6^. Decades of research have uncovered how an organism’s environment governs selection on display complexity by altering the transmission, perception, and fitness costs of multiple signals in a communication system^6–10^. By contrast, the evolutionary origins of complex displays remain elusive^11^. It is particularly difficult to envision how inter-signal interactions initially evolve, since at least some individuals must first produce adaptive signal combinations before selection can refine the signals’ integrated functions.

Moreover, display complexity often varies along continent-wide abiotic gradients, even among closely related species with similar sensory environments^12,13^. Yet, the mechanisms underlying these complexity gradients have never been experimentally tested. Resolving the origins and geographic diversity of complex communication requires empirically linking variation in the production of multiple signals to selection on complex display architectures, macroevolutionary gains in display complexity, and the consequences of display evolution for contemporary species distributions^5,14^.

Here, we test how environmental shifts affect the coordinated exaggeration and integrated functions of multiple signals, and whether changes in signal coordination can predict the environments where complex displays are most likely to evolve and persist. This hypothesis draws from longstanding theory regarding how phenotypic plasticity alters patterns of trait covariation and exposes new trait combinations to selection^15,16^. In our conceptual model, producing exaggerated levels of one signal type amplifies the receiver’s response to a second signal type (Figure 1A). Selection can only favor the coordination of these signals in environments that allow enough individuals to exaggerate both signals at the same time (*i.e.*, co- exaggeration; Figure 1B-C). This correlational selection should also be most efficient when the two signals show positive covariation, where individuals either exaggerate both or neither signal (Figure 1B-C). Environments that enable co-exaggeration and positive covariation between incipient signals could enable selection to initially favor complex displays and trigger the accumulation of additional signals over evolutionary time. On shorter time scales, environments that weaken signal coordination could disrupt effective communication. Such disruptions, if severe enough, may even preclude population viability^17,18^.

**Figure 1.**
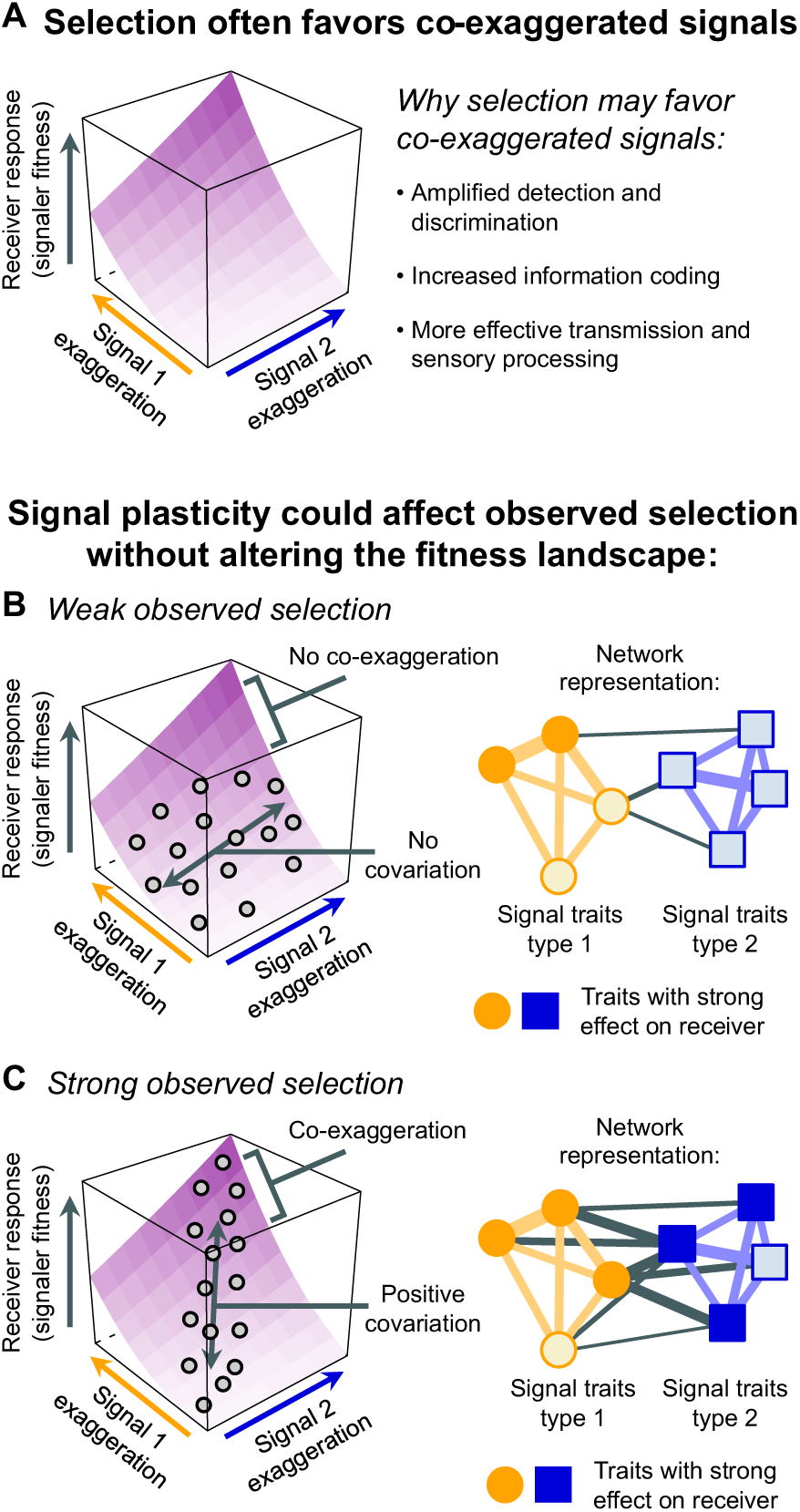
How signal coordination can enable selection for complex displays. **(A)** When the coordinated production of two signals amplifies the function of a display, producing exaggerated levels of both signals results in the strongest receiver response (i.e., increased signaler fitness; lighter regions of the surface plots^5,22^). **(B-C)** Plasticity in signal production can affect observed selection without changing the peak in the fitness landscape. For selection to efficiently act on the two integrated signals, enough individuals must co-exaggerate both signals simultaneously. Selection will also be more efficient if the signals show positive covariation, since individuals will either be exaggerating both or neither signal. **(B)** Environments that hinder signal co- exaggeration and positive covariation should reduce observed selection on both signals. For high-dimensional communication systems, reduced signal co-exaggeration and positive covariation can be represented by weaker between-type connections in a phenotype network. **(C)** Environments that promote signal co-exaggeration and positive covariation should strengthen observed selection on integrated signals. This is represented by stronger between-type connections in a phenotype network.

Signal combinations are particularly important in mating displays—an essential form of communication comprising some of the most exaggerated examples of display complexity^4^. Combining gestural or acoustic courtship behaviors with enlarged and pigmented morphological ornaments is a hallmark of sexual communication across animal taxa^4,19^. These two types of signals often interact in functional ways that favor the correlated evolution of highly complex courtship and exaggerated ornaments^20,21^. For example, highly exaggerated ornaments can enhance the detectability and attractiveness of courtship behavior, but producing ornaments alone often provides few benefits for mate attraction if the corresponding behaviors are hindered or absent^22–24^. Given that behavioral plasticity is often more dynamic than morphological plasticity^25^, environmental shifts could affect patterns of co-exaggeration, covariation, and the integrated functions of these two signal types^16^. Specifically, abiotic factors like temperature and water availability have widespread impacts on the production, function, and evolution of individual behavioral and morphological sexual signals^26–29^, but how climate shifts influence signal coordination and selection on display complexity has been overlooked^30^.

The hypothesis that plasticity in signal coordination shapes display complexity across climates makes predictions spanning behavioral, macroevolutionary, and community-level biogeographic processes. Specifically, (i) shifts in temperature and water availability that enable signal co-exaggeration and strengthen positive covariation between signal types should also promote mating display function (*i.e.,* increase the effect of display variation on mating success; Figure 1). (ii) Because these abiotic conditions increase the efficiency of selection on combined signals, species found in such environments should accumulate additional signal components and evolve increasingly complex displays. Moreover, because the functioning of sexual communication can be critical for successful establishment and persistence across species ranges^31–33^, (iii) climates that strengthen key inter-signal interactions should also harbor species assemblages with increased display complexity; and (iv) species that evolve complex displays should be more likely to persist in parts of their historical ranges where recent climate change is conducive to signal coordination. We tested these behavioral, macroevolutionary, and macroecological predictions using *Schizocosa* wolf spiders—an ideal system to examine how climate shapes display complexity, since *Schizocosa* species inhabit a range of climates^34^ and vary dramatically in both the number of courtship behaviors (gestures and vibrational songs) and the exaggeration of morphological signals (darkened and enlarged forelimb ornaments) that males use to attract mates^21,34,35^. Using a joint application of experimental, phylogenetic, and biogeographic approaches, we tested if changes in signal coordination predict changes in the complexity of species’ mating displays across historical North American climate gradients, along with species’ local persistence in response to recent climate change.

## Results

### Signal coordination enables selection for complex displays

We first experimentally tested how variation in temperature and water availability affected the coordinated production of behavioral and morphological signals. This experiment focused on *Schizocosa ocreata*, a highly ornamented species that also produces multiple gestural and substrate-borne vibrational courtship behaviors^35^ (Figure 2A). We collected subadult *S. ocreata* from a field site near St. Louis, MO USA and reared them to maturity under standardized laboratory conditions. After molting to adulthood, we manipulated adult water availability and temperatures during mating in a factorial design (cool-and-dry, cool-and-wet, hot-and-dry, hot-and-wet, each approximating shifts in the routine moisture and temperatures levels found in natural populations; Figure S1). We randomly paired males and females from each treatment combination for mating trials. For each trial, we measured 12 male sexual traits (eight behavioral and four morphological traits; Figure 2A). We then quantified the prevalence of co-exaggeration and the strength of positive covariation between behavioral and morphological signals in each of the four environmental treatments using phenotype networks—a systems-based approach that is emerging as a powerful tool for characterizing complex trait architectures^5^. In each phenotype network, “nodes” represent individual sexual traits that describe characteristics of a signal, and “edges” represent associations in the production of two sexual traits. Our phenotype networks revealed that hotter and wetter environments enabled more prevalent co-exaggeration between courtship and ornamentation (Figure 2B, Figure S5; Table S1). Hotter and wetter environments also led to stronger positive covariation between the two trait types (Figure 2C, Figure S5; Table S1).

**Figure 2.**
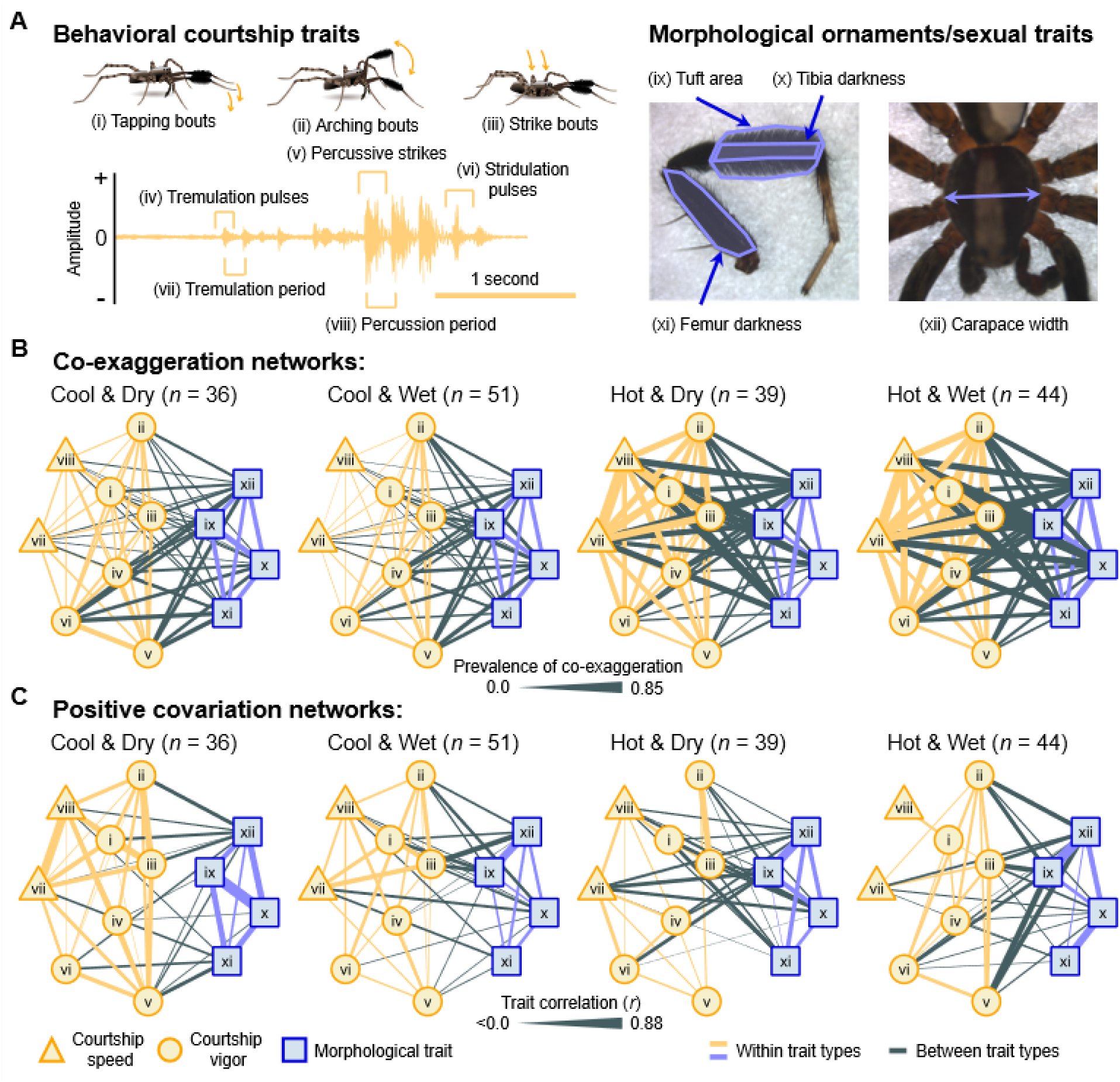
Courtship behaviors and morphological sexual traits are co-exaggerated and positively covary in hot and wet environments. **(A)** Behavioral and morphological sexual traits characterizing *Schizocosa ocreata* mating displays. Male *S. ocreata* use combinations of gestural (i-iii) and vibrational (iv-viii) courtship signals during courtship. The yellow oscillogram illustrates the relative amplitude and timing of vibrational signals. Gestural and vibrational signals are produced concurrently, with tapping and arching bouts performed during abdominal tremulation and cheliceral strike bouts acting as both a visual (gestural) and percussive (vibrational) signal. Male forelimb ornaments comprise enlarged tufts (ix) and dark pigmentation (x and xi). Male body size (carapace width; xii) is also assessed during female mate choice. **(B-C)** Phenotype networks illustrating the coordinated production of courtship speed traits (yellow triangles), courtship vigor traits (yellow circles), and morphological sexual traits (blue squares). Thicker edges between trait pairs indicate **(B)** more prevalent co-exaggeration and **(C)** stronger positive covariation.

We then used two complimentary approaches to test if changes in coordination between signal types influenced the function of mating displays, or the degree to which male display variation predicted mating success. First, we performed principal component analyses on the correlation matrices of the sexual traits in each of the four treatments (Table S2) and examined how variation along each principal component affected mating success. Consistent with our predictions, PC1 in the hot-and-wet treatment had the strongest effect on mating success compared to all principal components in other treatments (hot-and-wet PC1 *β* = 2.03, 95%CI: 0.77 to 3.95, *P* < 0.001; Figure 3; Table S3), and this PC corresponded to covariation in both behavioral and morphological traits (Figure 3; Table S2). Second, we tested how rates of signal co-exaggeration differed between males that successfully mated and males that did not mate.

**Figure 3.**
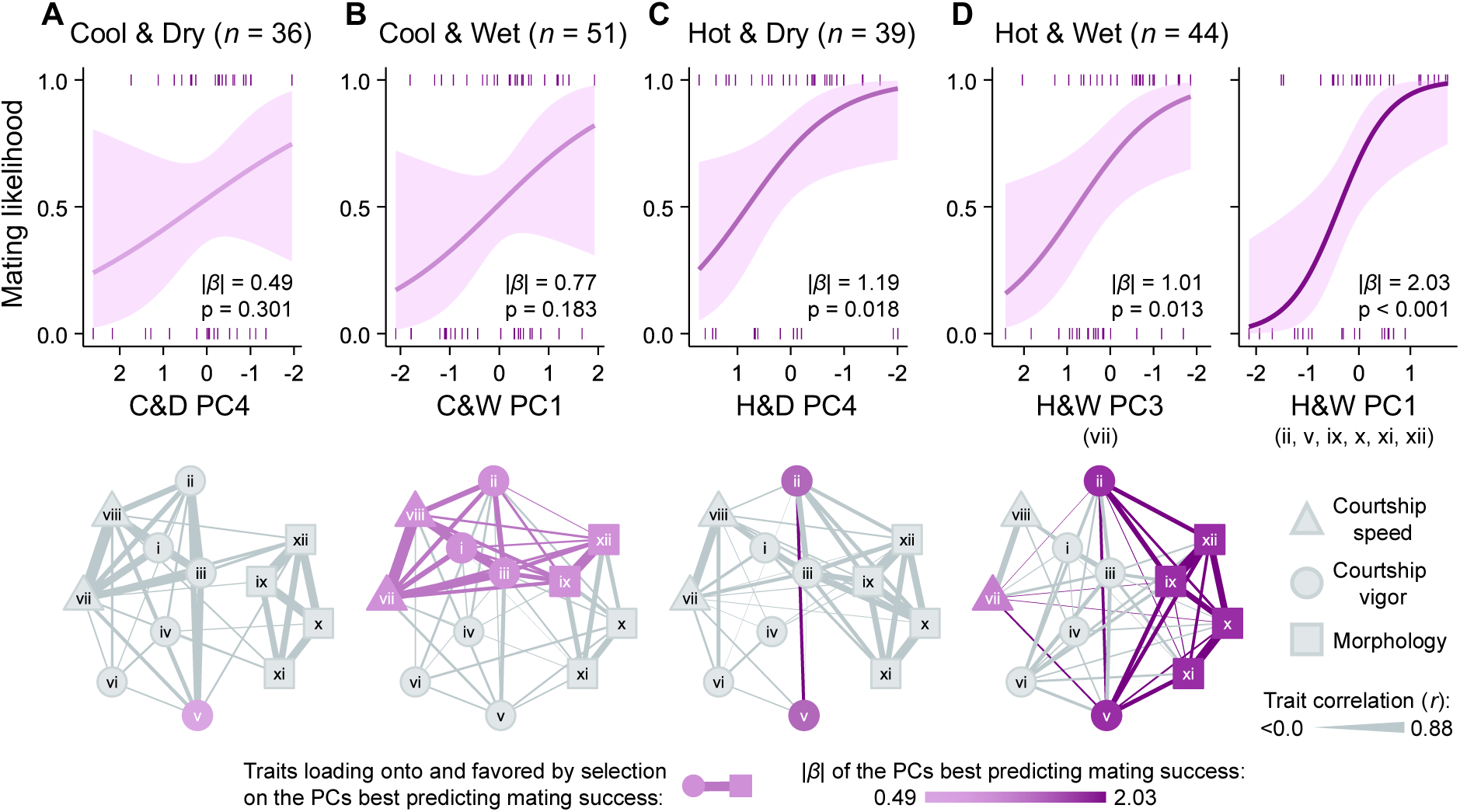
Signal covariation enhances the function of complex mating displays for *Schizocosa ocreata* in hot and wet environments. Principal components representing covariation among sexual traits had stronger effects on mating success in the hot & wet treatment. The lines and confidence bands show GLM estimated fits and 95% confidence intervals for the principal components that best predicted mating success in the cool & dry treatment **(A)**, cool & wet treatment **(B)**, hot & dry treatment **(C)**, and hot & wet treatment **(D)**. Two principal components significantly affected mating success in the hot & wet treatment, and both are shown with the traits corresponding to each PC in parentheses below the x axes. Phenotype networks below each mating effect plot show the traits that load strongly onto (magnitude of loading factor > 0.4) and are favored by estimated selection on the PC that best predicted mating success in each treatment (purple traits and edges). The darkness of the purple nodes, edges, and model fits indicates the magnitude of the standardized effect estimate on mating success (|*β*|). Courtship speed traits are shown in triangles, courtship vigor traits are shown in circles, and morphological sexual traits are in squares. Thicker edges between traits indicate stronger positive covariation between trait pairs.

Mated males were significantly more likely to have co-exaggerated behavioral and morphological traits only in the two hot treatments, with the strongest effect of co-exaggerating behavioral and morphological traits on mating success in the hot-and-wet treatment (Figure S6). Hotter and wetter conditions thus increased the coordinated production of behavioral and morphological signals, which in turn enhanced the integrated functions of those signals for mating success.

### Changes in signal coordination predict the evolution of display complexity

The findings from our mating trials showed that exposure to hotter and wetter environments can enable selection to favor combinations of behavioral and morphological signals. If hotter and wetter climates also promote functional interactions between existing and incipient signals, then taxa exposed to these climates should accumulate additional courtship behaviors alongside more exaggerated ornaments over evolutionary time. To test this prediction, we compiled the total number of courtship behaviors^21,35–50^ and standardized ornament exaggeration scores^34^ for 42 taxa represented in a recent *Schizocosa* phylogeny^34^ (Figure 4A). We then extracted historical climate data^51^ for each taxon in our database (Figure 4A). Phylogenetic logistic regression showed that taxa are more likely to evolve both additional courtship behaviors and greater ornament exaggeration in hotter (*β* = 1.21 ± 0.57, 95%CI: 0.26 to 2.88; Figure 4B; Table S5) and wetter climates (*β* = 0.71 ± 0.39, 95%CI: 0.01 to 1.86; Figure 4C; Table S5). A supplementary analysis treating display complexity as a continuous, composite score of behavioral complexity and ornament exaggeration also showed that lineages in hotter and wetter regions have evolved more complex displays (Figure S7; Table S7).

**Figure 4.**
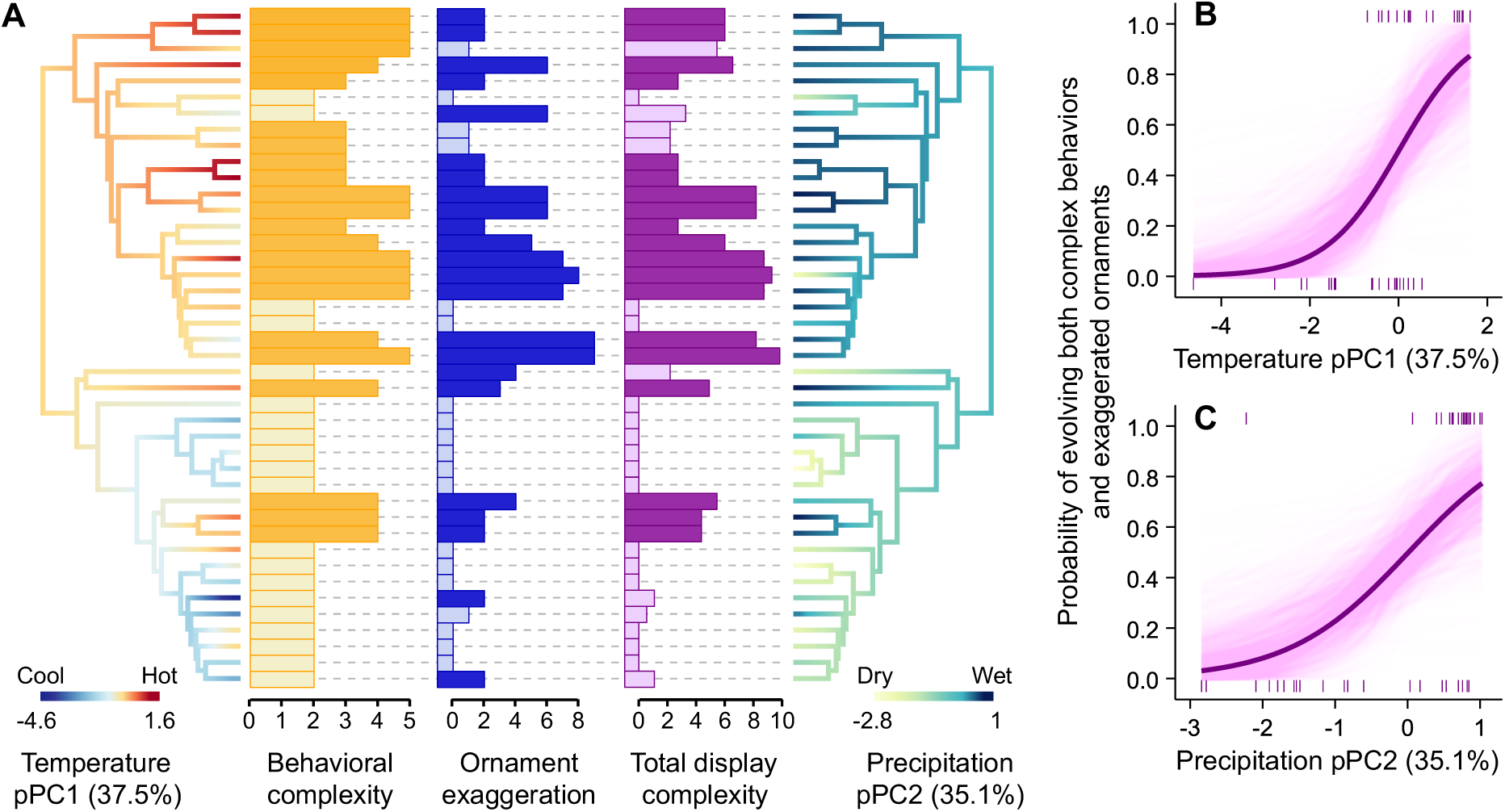
Species in hotter and wetter climates evolve increased display complexity. **(A)** *Schizocosa* phylogeny illustrating among-taxon covariation in behavioral complexity (total number of courtship behaviors; yellow), ornament exaggeration scores (blue), total display complexity scores (purple), and the phylogenetic PCA scores for bioclimatic variables. Darkened bars for behavior and ornament traits indicate values greater than or equal to the mean values among taxa, and darkened bars for total display complexity indicate taxa that have evolved at least the median values for both behavioral complexity and ornament exaggeration. For pPC1, negative values (bluer branches) indicate colder climates and positive values (redder branches) indicate hotter climates. For pPC2, negative values (yellower branches) indicate drier climates and positive values (bluer branches) indicate wetter climates. The ancestral states of the climate PCs were used only for visualization and were not in a formal analysis. **(B)** Taxa are more likely to evolve both high behavioral complexity and high ornament exaggeration in hotter climates. **(C)** Taxa are also more likely to evolve both high behavioral complexity and high ornament exaggeration in wetter climates. Points in **B** and **C** indicate individual taxa (*n* = 42), bolded lines are model estimates, and transparent lines show estimates from 1000 bootstrapped phylogenetic logistic regressions.

To evaluate whether the observed evolutionary gradients in display complexity more likely emerged from correlational selection favoring adaptive signal combinations, or selection acting independently on behavioral and morphological signals, we performed a phylogenetic path analysis. The best supported path model showed that hotter and wetter climates indirectly affected ornament exaggeration (Table S8). Additional courtship behaviors are more likely to evolve in hotter (*β* = 1.98 *±* 0.94, 95%CI: 0.57 to 2.81) and wetter climates (*β* = 1.45 *±* 0.54, 95%CI: 0.46 to 2.65), which then enables additional behaviors to co-adapt with increased ornament exaggeration (*β* = 3.00 *±* 0.82, 95%CI: 1.49 to 4.66; Figure S8), rather than each signal type evolving independently. Collectively, our phylogenetic comparative results demonstrate that the evolution of complex displays is biased to environments that strengthen signal coordination and integrated signal functions.

### Changes in signal coordination predict gradients in assemblage composition

Our continent- wide phylogenetic analyses indicated that hotter and wetter climates have driven the correlated evolution of complex courtship and exaggerated ornaments in parallel to how these climates enhance the integrated function of these signals. However, at finer spatial scales, whether ecological communities comprise species with complex versus simple displays depends not only on *in situ* display evolution, but also on species filtering^52^. Just as reduced fertility in extreme climates can limit species’ geographic ranges^53^, climates that hinder key display functions should also limit establishment by species that depend on complex displays for reproduction. Other assemblage-level processes could also provide alternative explanations for why display complexity increases in hotter and wetter climates. For instance, complexity may instead evolve because it enhances species recognition in tropical and subtropical regions where closely related species often coexist^54,55^.

We therefore used a spatially explicit analysis to test how an area’s climate affected the mean total display complexity (a combined score of behavioral complexity and ornament exaggeration) within assemblages of *Schizocosa* species throughout North America. We quantified 549 assemblages (Figure 5A) using 7,229 *Schizocosa* occurrence records^56^ and climate data during the mating season^51^. Spatially explicit generalized least squares confirmed that assemblages comprise species with more complex displays in hotter (temperature PC2 *β* = 0.32 ± 0.06 SE, 95%CI: 0.19 to 0.44, *P* < 0.001) and wetter climates (precipitation PC1 *β* = 0.51 ± 0.08 SE, 95%CI: 0.36 to 0.66, *P* < 0.001, Figure 5B, Table S11). Moreover, species richness in the assemblages did not affect assemblage-mean display complexity (*β* = 0.04 ± 0.04 SE, 95%CI: -0.04 to 0.12, *P* = 0.258, Table S11). Gradients in display complexity are therefore not associated with predicted selection favoring complex species recognition signals—a commonly invoked mechanism driving display complexity in more tropical areas^54,55^. Assemblage-level variation in complexity more likely emerges from species filtering due to impaired display function in cool and dry climates or enhanced display function in hot and wet climates.

**Figure 5.**
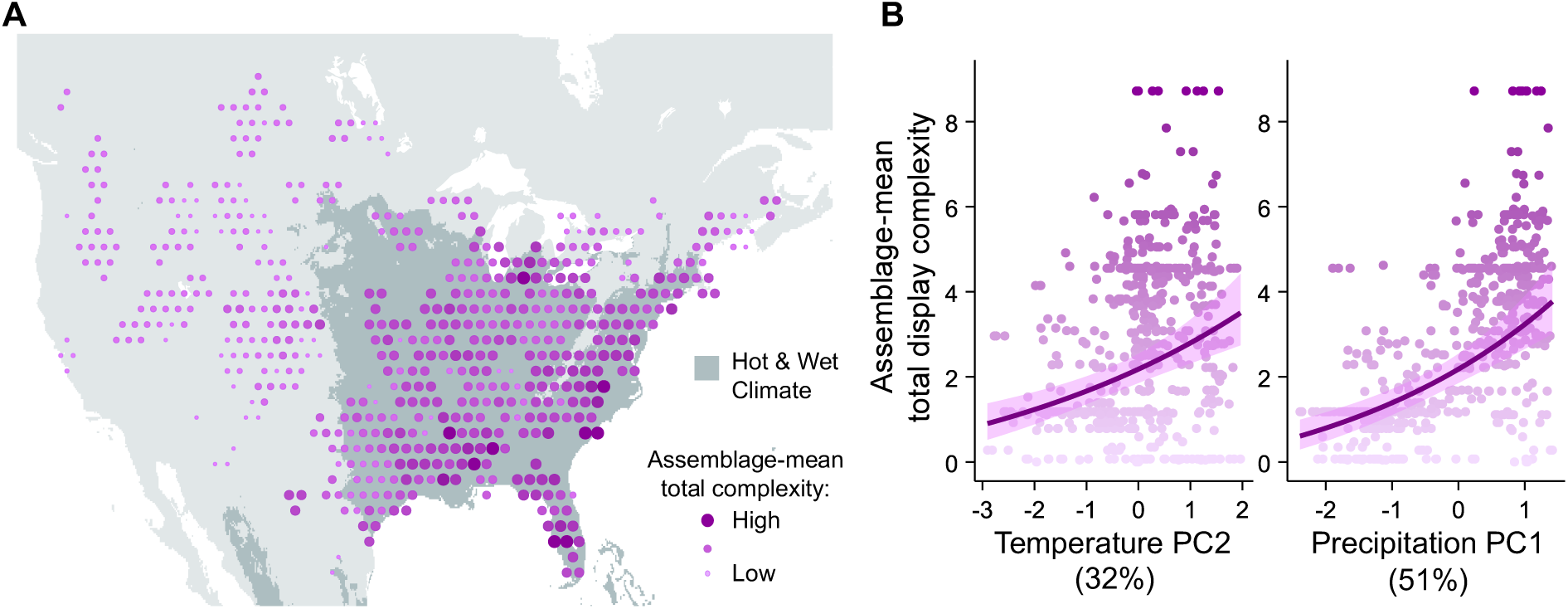
Assemblages in hotter and wetter climates comprise species with increased display complexity. Points in all panels show sliding-window assemblages (*n* = 549) grouping GBIF occurrence records of *Schizocosa* species. Darker and larger points indicate assemblages with a higher mean total display complexity among species in the assemblage. Shaded geographic regions in **(A)** are tropical, subtropical, or temperate with hot summers and no dry season (Köppen-Geiger designations: Af, Am, Aw, Cfa, Cwa, and Dfa^68^). Positive PC2 values indicate hotter climates and positive PC1 values indicate wetter climates. In **(B)**, lines are model estimates and bands are 95% confidence intervals from the spatially explicit GLS with the best supported spatial correlation structure.

### Complex display evolution predicts persistence following recent climate change

Contemporary climate change is also rapidly altering species’ ranges^57^, allowing us to further test if species persistence within their historical ranges is associated with the predicted effects of climate change on complex display function. Recent evidence suggests that animal populations are especially vulnerable to extinction in geographic areas where climate change impedes crucial reproductive processes^32,58^. *Schizocosa* species that depend on complex displays during reproduction should similarly persist less often in areas that are getting cooler and drier, as these conditions should disrupt signal coordination and function. Because effective sexual communication can also provide extra buffers against extinction in changing environments^33,59^, species with complex displays may even persist more often than other species in increasingly hot and wet climates.

We tested how recent climate change across North America has shaped the local persistence of *Schizocosa* species that have evolved both high behavioral complexity and high ornament exaggeration. We first created a dataset of 75 geographic areas (2° latitude grid cells) spanning the historical ranges of 12 abundant species^60^. For each area, we determined whether species that were previously observed in the 1970s-90s have persisted to the present, and calculated the change in temperature and precipitation within each area between these two periods^51^. We used generalized linear mixed models^61^ that accounted for spatial and phylogenetic non-independence to test for relationships between species’ display complexity and persistence likelihood^32^ (Table S13). Species with complex displays are more likely to persist in areas that are becoming warmer (*β* = 1.26 *±* 0.43, 95%CIs: 0.42 to 2.10; Figure 6A; Table S14) and wetter (*β* = 0.74 *±* 0.34, 95%CIs: 0.08 to 1.41; Figure 6B; Table S14), while the persistence of species with simple displays is not significantly affected by recent climate change (Figure 6; Table S13). These patterns of recent species persistence recapitulate our experimental and comparative results. Even after species evolve complex displays, their subsequent persistence is limited to environments that also promote the integrated functions of multiple signals.

**Figure 6.**
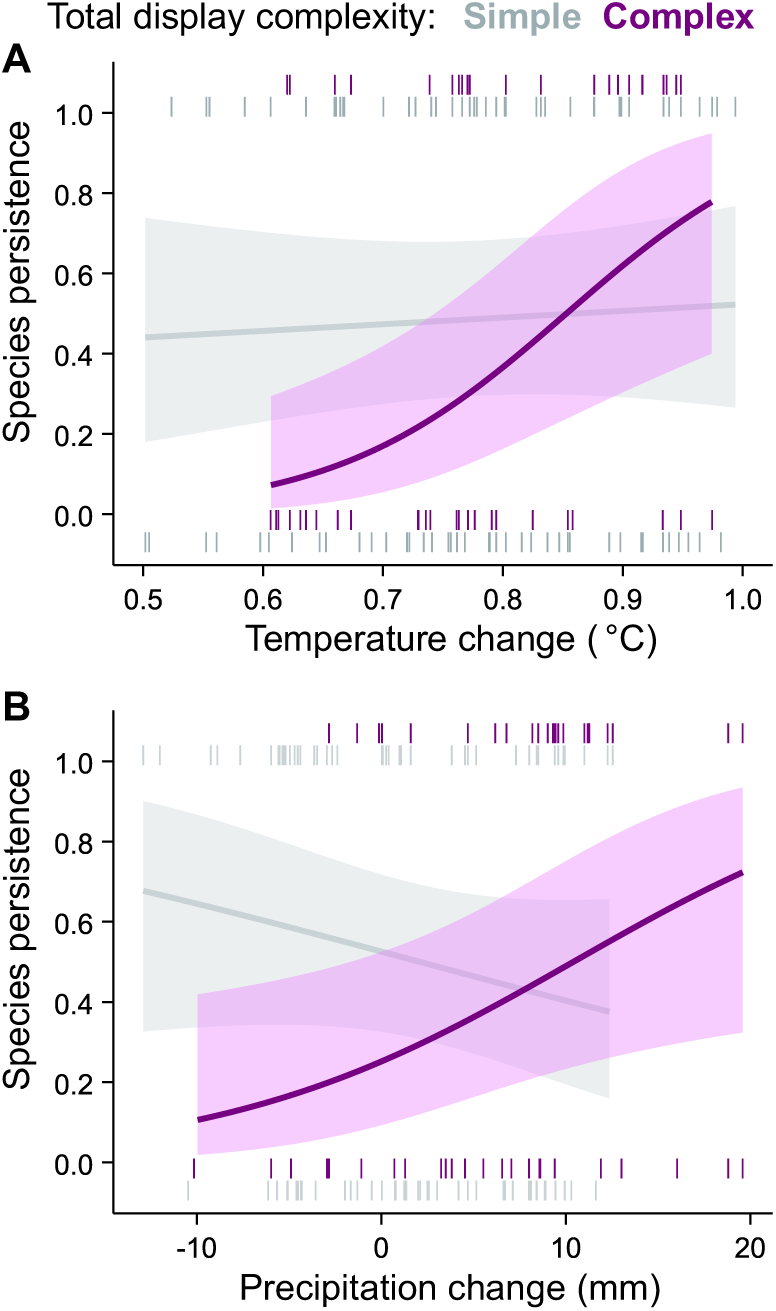
Species with complex displays are more likely to persist in areas that recently became hotter and wetter. *Schizocosa* species with complex displays (species that have evolved at least the median number of courtship components and ornament exaggeration scores among species in the phylogenetic analysis; purple points and lines) are more likely to persist in areas exposed to more severe climate warming **(A)** and increasing precipitation **(B)** from the 1970s-90s to today compared to species with relatively simple displays (grey points and lines. Points indicate individual species persistence measurements (n = 153) among the 12 species (five species with complex displays and seven species with simple displays) included in the analysis of 75 grid cells. Lines are model estimates, and bands are 95% confidence intervals from GLMMs.

## Discussion

Joint assessment of our experimental, comparative, and biogeographic findings (a powerful framework for evaluating causal ecological and evolutionary hypotheses^62^) suggests that changes in signal coordination are likely important for the repeated evolution of complex displays in hotter and wetter climates. Species with complex courtship and exaggerated ornaments were also more likely to persist in areas experiencing more severe climate warming and intensified precipitation over the last 50 years. Environmental effects on the production of multiple functionally integrated signals are therefore aligned with how display evolution predicts population viability, potentially further shaping contemporary geographic variation in display complexity.

Signals used for mate attraction are crucial for population-level fitness in altered environments because they can facilitate mate localization, species recognition, and mate choice favoring individuals with locally adaptive alleles^31,33,59^. Understanding how sexual signals have adapted to past climates can therefore reveal how populations may evolve or go extinct in the wake of future climate change^28,32,63^. However, studies to date on how climate change affects sexual communication have overwhelmingly explored responses in one signal at a time^30^, overlooking critical changes in integrated signals that we demonstrated here. Our results also emphasize that researchers should examine how multiple climate stressors simultaneously influence sexual communication to understand how climate impacts population fitness ^64^. Even though an organism’s sensitivity to temperature is intricately connected to hydration state ^65^, only a handful of studies examine the effects of altered water availability on sexual trait function (e.g., studies in already water-limited species^66,67^) compared to the rapidly accumulating evidence that warming influences sexual communication^26–28^. Our experimental findings suggest that decreased or more stochastic precipitation could hinder the function of complex displays even in habitats that show no temperature change. Many animals in the Nearctic, however, are likely to experience moderate increases in both temperature and precipitation in the next century, especially at typically cooler high latitudes^68^. Most high-latitude *Schizocosa* currently possess relatively simple mating displays, but may begin to accumulate additional courtship behaviors or more exaggerated ornaments if future climates strengthen covariation among incipient signals. By contrast, the decreased precipitation forecasted in the American Southwest and Mexico in the next century^68^ may compromise the functions of *Schizocosa* displays for mate attraction^28^. If individuals cannot seek out suitable temperatures or the water resources required for display function^69^, our results suggest that climate-induced breakdowns in sexual communication may threaten population persistence.

Considering how the environment influences signal coordination also brings to light new ways in which communication systems should co-adapt with other aspects of an organism’s phenotype. Biologists are increasingly recognizing that signals co-adapt with receivers’ sensory, neural, and cognitive biology in response to selection on these traits in non-signaling contexts^8,70^. For example, differences in the light environments among surfperch species favor the evolution of distinct photoreceptor sensitivities for prey detection in females, and male sexual coloration has diversified to match these differences in female visual systems^71^. Our results emphasize that signals should also co-adapt with signaler traits that dictate the environmental sensitivity of signal production^28,72^. Signal coordination likely changes across environments because distinct physiological processes affect the production of different types of signals (gestures, sounds, coloration, and structures), and these physiological processes themselves differ in plasticity^28,73^. Once multiple signals begin to covary due to plasticity in a new environment, the developmental integration of multiple physiological processes may evolve to further maximize signal co- exaggeration and display function (i.e., ‘genetic accommodation’)^15,16,73^. Future investigations into how plasticity-first evolution shapes divergence in complex communication will therefore also advance our understanding of eco-physiological adaptation^16,74^.

The evolution of communication is fundamentally tied to variation in the signaling environment^7^. Yet, how the environment affects the production of multiple interdependent signals is an underappreciated process that should operate in tandem with environmental effects on signal transmission, detection, and perception to guide divergence in communication systems^75^. Our results demonstrate how changes in signal coordination regulate the environments where selection can efficiently act on multiple display components, which has likely biased the environments where display complexity has accumulated over evolutionary time and the geographic regions where species with complex displays are most likely to persist. These findings unify foundational theory regarding how multivariate plasticity affects selection on complex features^15,16^ with a modern systems-inspired theory of animal communication that emphasizes inter-signal interactions^5,6^. Overall, considering environmental effects on signal coordination sheds light on the enigmatic origins of complex communication, revealing a novel association between signal production and continent-wide patterns of biogeographic diversity.

## Methods

### Signal coordination enables selection for complex displays

We performed a laboratory mating experiment using *Schizocosa ocreata* to determine if and how climate conditions affect the coordinated production of behavioral and morphological sexual traits, and whether climates that strengthen coordination between the two signal types also caused displays to have stronger effects on male mating success. We collected subadult *S. ocreata* on April 27-May 1, 2021, at a field site near St. Louis, Missouri, USA (Lat: 38.554291; Long: -90.369585). Spiders were reared to adulthood under laboratory conditions in individual 6cm-6cm-8cm clear plastic containers (AMAC 760C boxes) that were lined with vinyl screens on two inside surfaces. All individuals were kept at an ambient temperature of 23°C, experienced a 12hr:12hr light:dark cycle, and were fed one 3mm cricket (*Gryllodes sigillatus*) every three days. Before molting to adulthood, we provided all individuals with continuous access to water by inserting a cotton wick through the container floors that absorbed water from a reservoir beneath the containers.

#### Water and temperature treatments

We manipulated adult water availability and mating trial temperatures in a full factorial design. Two weeks prior to the mating trials, we randomly assigned adult males and females to one of two water availability treatments. We maintained watering treatments through adulthood and selected males and females from the same treatment for all mating trials. Individuals in the ‘wet’ treatment received a new cotton wick through the watering hole in their container floors and had continuous access to water. For individuals in the ‘dry’ treatment, we inserted new cotton wicks through the floors of the containers but did not connect them with water reservoirs. Instead, we placed a 25µL drop of water on the container floors every three days during regular feeding and cage cleaning. We assessed the ecological relevance of the watering treatments by comparing differences in hydration state (body water content) between treatments to variation among field-active adults^76^ (Supplemental Information). Providing continuous water access in the laboratory led to hydration states matching the most hydrated individuals in the field (Figure S1A). Individuals in the dry treatment had lower hydration states, but not as low as the most dehydrated individuals in the field (Figure S1A).

However, restricting water access beyond three days led to high mortality, indicating that our design maximized the possible differences in water availability under controlled laboratory conditions. Overall, our watering treatments are representative of natural variation in water access/hydration, but it is possible that other conditions in nature allow spiders to survive at slightly lower hydration states (*e.g.,* diet changes^77^).

We ran each mating trial at one of five target temperatures (15, 20, 25, 30, or 35°C) that span the range typically experienced in the field (Figure S1B). Courtship and mating were rare outside this temperature range in preliminary trials. We used these five target temperatures to explore nonlinear changes in mating behavior across a continuous temperature gradient ^26,29^ using thin-plate splines (‘*fields*’^78^) in R v4.2.0^79^. See Supplemental Information for detailed methods and results regarding nonlinear temperature responses. The effect of each sexual trait on copulation success always changed monotonically with increased temperature, and often showed an inflection point as temperatures surpassed the median testing temperature, 25.3°C ( Figure S3 and Figure S4). We therefore categorized testing temperatures as either ‘hot’ (greater than 25.3°C) or ‘cold’ (less than or equal to 25.3°C) for analyses reported in the main text due to sample size requirements for multivariate analyses. This categorization led to four environmental treatment groups: cold-and-dry, cold-and-wet, hot-and-dry, and hot-and-wet. We held temperatures constant throughout the trials and measured exact arena temperatures with a type-K thermometer (Fisher Scientific Traceable 91210-07).

#### Mating trials

We began mating trials three weeks after spiders reached adulthood, when female receptivity peaks^80^. We conducted mating trials between randomly paired males and females within custom incubators to manipulate trial temperature (Supplemental Information). Inside the incubators, we placed a 20cm-diameter, 5cm-tall circular acetate mating arena that was lined on the outside with brown construction paper. We also lined the floor of the arena with brown construction paper, which resembles the color and vibrational properties of the leaf litter used during courtship in the field^45^. We first acclimated the female within the arena for 20 minutes. Males were simultaneously acclimated within the testing incubator but in an isolated container. This acclimation period allowed females to lay down silk on the arena floor, which reliably elicits male courtship^45^. We then placed a 5cm-diameter transparent acetate barrier around the female, moved her to the center of the arena, and introduced the male outside of the barrier to prevent instantaneous mating or cannibalism. We removed the barrier two minutes after the onset of male courtship (or after five minutes if the male did not court) and allowed the male and female to interact freely. We ended the trial when mating occurred or five minutes after removing the barrier if mating did not occur. Previous work suggests that this trial duration is sufficient to assess mating decisions^81,82^, and when mating occurred, the mean latency to mate after the barrier was removed was 1.26 minutes (SE ± 0.34). We recorded videos for all trials using a Sony 4K high-definition camera (Model FDR-AX53). Audio files of substrate-borne vibrational courtship were recorded in the program Audacity v3.4.1 using a PCB Piezotronics accelerometer (Model 352A24 with signal conditioner Model 480E09) and a Roland Duo-Capture USB audio interface (Model UA-11-MK2). Out of 183 trials, males did not perform enough courtship to measure all behavioral traits in 13 trials (4 cool-and-dry; 1 cool-and-wet; 5 hot-and-dry; 3 hot-and-wet). We therefore included data from 170 trials in our final analyses (cool-and-dry: 36; cool-and-wet: 51; hot-and-dry: 39; hot-and-wet: 44).

#### Sexual trait measurements

For males in each mating trial, we measured eight behavioral courtship traits (Figure 2A) that resemble putative sexual traits used to describe mating signal characteristics in other *Schizocosa* species^29,36,48^. We measured the total number of (i) leg tapping bouts, (ii) leg arching bouts, and (iii) cheliceral strike bouts produced during the first minute of male courtship. These components involve both gestural and vibrational signals. Leg tapping and leg arching bouts are accompanied by simultaneous abdominal tremulation, while cheliceral strikes are a single behavior that produces both a gestural and a percussive signal. These three main components are often (but not always) produced in stereotyped displays comprising either a leg tapping bout or a leg arching bout, followed by cheliceral strikes, followed by pedipalp stridulation (Figure 2A). Within stereotyped displays, we measured the mean number of (iv) tremulation pulses, (v) percussive strikes, and (vi) stridulation pulses, along with the mean period between (vii) tremulation pulses and (viii) percussive strikes. Traits iv-viii were averaged across the final three ordered displays that occurred in the first minute of courtship.

We also measured four male morphological traits that are involved in mate attraction (Figure 2A)^46,80^ using a Leica dissecting microscope (Model M205 C) and ImageJ^83^. All images of male morphological traits were captured on the same day to maintain consistent lighting conditions. We measured (ix) forelimb ornament size from the area of a ten-sided polygon outlining the total ornamental tuft, along with (x) the darkness of the forelimb tibia and (xi) the darkness of the forelimb femur by measuring the mean gray value (on a scale from 0 to 255, where 0 is black and 255 is white). We estimated male body size by measuring (xii) the width of the carapace along its widest point perpendicular to its longest axis.

#### Phenotype networks

We constructed phenotype networks to analyze patterns of co-exaggeration and positive covariation among the 12 male sexual traits^36,84^. We constructed networks separately for each of the four environmental treatment groups and visualized networks using the R package ‘*qgraph*’^85^. The nodes in these networks represent individual sexual traits, and the edges represent trait associations (*i.e.*, co-exaggeration prevalence or positive covariation). To characterize patterns of trait co-exaggeration, we first determined whether or not males produced exaggerated levels of each trait (greater than the median trait value across all treatments; 1 = exaggerated, 0 = unexaggerated). Then, for each pair of traits in the network, we computed the prevalence of co-exaggeration from proportion of males in the environmental treatment that had exaggerated levels of both traits simultaneously. To characterize patterns of positive trait covariation, we first created a Pearson’s correlation matrix for the traits in each environmental treatment and then set all negative trait correlations to zero.

We tested how environmental treatments affected patterns of co-exaggeration and positive covariation between behavioral and morphological sexual traits by assessing changes in two complimentary indices. First, we calculated the average weight of edges connecting behavioral and morphological traits in each treatment. We calculated 95% confidence intervals around these average between-type edge weights from 1000 bootstrapped estimates, resampling the between-type edges with replacement for each bootstrap iteration. Second, we computed the assortativity of edges within trait types (*r*_d_) with the R package ‘*assortnet*’^86^. We calculated standard errors for *r*_d_ using jackknife resampling with *assortnet* because the *r*_d_ statistic is not normally distributed. Here, *r*_d_ describes the relative magnitude of edges that were within versus between behavioral and morphological traits^84^. If one environmental treatment increased relative co-exaggeration or positive covariation between behavioral and morphological traits compared to other treatments, the network from this treatment will have a smaller *r*_d_ value.

#### Display function for mating success

We used two complimentary approaches to test if behavioral and morphological sexual traits had stronger effects on mating success in environments where they were strongly coordinated. For the first approach, we performed separate principal components analyses on the sexual trait correlation matrices in each of the four environmental treatments (‘*psych*’^87^). For each treatment, we extracted scores from five principal components representing the major axes of phenotypic covariation among the 12 male sexual traits (Table S2)^36,84^. We tested how variation along these principal components affected mating success in each treatment using binomial generalized linear models with whether the male-female pair copulated as the response and the five principal components as explanatory variables. Because mating temperatures varied within the binned temperature treatments (see *water and temperature treatments*), we included trial temperature as a covariate in all models. We initially included female age after maturity as a covariate^80^ but removed it from all models because it was never significant. All variables were z-transformed to standardize parameter estimates^88^. If coordination between behavioral and morphological traits enhanced signal function, then principal components in treatments with the greatest average between-type edge weights and lowest phenotype network *r*_d_ value should have stronger standardized effect magnitudes on mating success—particularly principal components that correspond to variation in both behavioral and morphological traits.

For the second approach, we directly tested how patterns of signal co-exaggeration predicted mating success. We created co-exaggeration networks for mated and non-mated males in each treatment using the methods described above and subtracted the edge weights of the non- mated networks from the mated networks. We then used permutation tests to assess whether differences in co-exaggeration between mated and non-mated males where greater than expected by chance. We permuted mating status across all males in the treatment 1000 times, and for each trait pair, compared the observed co-exaggeration difference to the null distribution of co- exaggeration difference between mated and non-mated males (Figure S6).

### Changes in signal coordination predict the evolution of display complexity

We leveraged a recently published *Schizocosa* phylogeny^34^ to test how temperature and precipitation have shaped the evolution of display complexity across the geographic extents of the genus. The phylogeny produced by Starrett and colleagues^34^ reveals that multiple historically recognized species groups are not monophyletic. Furthermore, intraspecific geographic variation in sexual signals is pervasive throughout the genus^34,38,43^. We therefore chose to examine patterns of trait evolution among the *Schizocosa* populations represented in the phylogeny by Starrett and colleagues^34^, which in total comprise 22 historically recognized species. We pruned the phylogeny to 42 taxonomically distinct populations, determined the total number of distinct courtship behaviors used by each taxon from published courtship descriptions^21,35–50^, and combined these data with the standardized ornament exaggeration scores reported by Starrett and colleagues^34^. See Supplemental Information for further details on taxon selection and characterizing sexual traits. Overall, this sexual trait database represents both intraspecific and interspecific variation among 42 distinct taxa, with each taxon being a single population from one of 22 historically recognized species. We then quantified each taxon’s total display complexity using two approaches. First, we identified whether each taxon did or did not evolve both high behavioral complexity and high ornament exaggeration by determining whether each trait was greater than or equal to the median trait value across all taxa (darkened bars in Figure 4A), resulting in 19 taxa with “complex” displays and 23 taxa with “simple” displays (see Table S15 for each taxon’s display description and complexity designation). Second, we rescaled the behavior and ornament traits so that they varied between zero and five across all taxa and then added these rescaled scores together. This produced a continuous total display complexity score that was equally weighted by variation in behavior and ornamentation.

#### Climate data

At the geographic coordinates provided for each taxon in the phylogeny^34^, we extracted four bioclimatic variables from WorldClim 2.0^51^: (i) mean temperature of the wettest quarter; (ii) maximum temperature of the warmest month; (iii) precipitation of the wettest quarter; and (iv) precipitation of the driest month. Both precipitation variables were log_e_- transformed before analysis. We selected these variables *a priori* to represent the conditions likely experienced during the mating season (typically the wettest annual quarter, or March-July^21,35–47^) and the most extreme conditions that could be experienced by each taxon throughout their life cycle. We reduced dimensionality among the bioclimatic variables using the first two axes from a phylogenetic principal component analysis (‘*phytools*’^89^). These axes together explained 72.1% of the climatic variation, with the first axis (pPC1, 37.0%) showing a strong positive relationship with temperature and the second axis (pPC2, 35.1%) showing a strong positive relationship with precipitation (Table S4).

#### Phylogenetic comparative methods

We used a phylogenetic logistic regression to test if taxa are more likely to evolve both high courtship complexity and high ornament exaggeration in hotter and wetter climates (‘*phylolm*’^90^). We included whether each taxon evolved both high courtship complexity and high ornament exaggeration as the response variable (see above and Supplemental Information) and temperature (pPC1), precipitation (pPC2), and their interaction as the explanatory variables (Ornstein-Uhlenbeck *α* = 3.7). We evaluated the significance of each effect by testing if the confidence intervals from 1000 bootstrapped parameter estimates overlapped zero. A supplementary analysis using the continuous total display complexity trait as the response in a phylogenetic generalized least squares model (‘*phytools*’^89^) showed similar results (Figure S7; Supplemental Information). We then used phylogenetic path analysis (‘*phylopath*’^91^; see Supplemental Information) to compare six hypothesized causal path models that varied in the inclusion of direct and indirect relationships among behavioral courtship complexity, ornament exaggeration score, temperature (pPC1), and precipitation (pPC2). We compared support among models by assessing CICc^91^ and performed model averaging when two or more path models were equally supported (ΔCICc < 2).

### Changes in signal coordination predict gradients in assemblage composition

We used an assemblage-level analysis that tested how the mean total display complexity among species in each assemblage related to the area’s climate. To quantify local *Schizocosa* assemblages, we gathered 7,229 georeferenced occurrence records from the Global Biodiversity Information Facility^56^ for the 20 historically recognized species that had available courtship, ornamentation, and occurrence data. We then grouped these occurrence records into 549 overlapping assemblages using a sliding-window approach^92,93^. A sliding-window approach was preferred because it minimizes assumptions about assemblage boundaries when using aggregated biodiversity data; however, a method using non-overlapping grid cells produced consistent results (Supplemental Information; Tables S9, S10, S11). Although the sliding-window approach inherently generates spatial autocorrelation in assemblage-mean phenotypes, this spatial structure was accounted for in our analysis (see below). We first generated a grid of 1° latitude-wide hexagonal cells across the geographic extents of the occurrence records^94^. Around each hexagonal cell centroid, we created a circular buffer zone with a 2° latitude diameter (approximately 31000 square km)^95^. These buffer zones represent a series of assemblage boundaries that each overlap approximately half the diameter of its neighboring zone. For each zone, we determined the mean display complexity score among all species observed in the zone. For species that were represented by multiple taxa in our phylogenetic analyses^34^, we used the mean complexity score among every taxon associated with that species. We also quantified species richness and the total number of observations included in each zone. We only included zones where at least two species were observed^92^. Following the justification outlined in our phylogenetic analyses, we determined the mean values of four bioclimatic variables across the entire area of each zone^51^: (i) mean temperature of the wettest quarter; (ii) maximum temperature of the warmest month; (iii) precipitation of the wettest quarter; and (iv) precipitation of the driest month. Both precipitation variables were log_e_-transformed before analysis. A principal components analysis on these bioclimatic variables yielded two components that collectively explained 83% of the variation in the dataset. We varimax-rotated these components to improve the interpretability of our results^87^, resulting in one component that showed a strong positive association with temperature (PC2 32%) and another component that showed a strong positive association with precipitation (PC1 51%; Table S9).

We used spatially explicit generalized least squares models (spatial GLS; ‘*nlme*’^96^) to assess the relative importance of temperature, precipitation, and species richness for shaping variation in display complexity among assemblages. We fit the log_e_-transformed(+1) assemblage-mean display complexity score as the response, with effects of temperature (PC2), precipitation (PC1), the interaction between PC2 and PC1, and log_e_-transformed species richness. To account for variation in sampling effort, we included the log_e_-transformed total number of observations in the buffer zone as covariate. All variables were also z-transformed to standardize regression coefficients^88^. We used a rational quadratic correlation structure in our final models because they received better support than models with other possible correlation structures (Table S10).

We also evaluated the sensitivity of our results to potential pseudo replication due to repeated species co-occurrences by comparing our observed effects to null distributions of expected test statistics^97^. We randomized the display complexity scores among the 20 species in our spatial dataset, recomputed the assemblage-mean values for each assemblage zone, and reconstructed 100 spatial GLS models to generate null distributions of 100 effect estimates ( *β*) for temperature (PC2), precipitation (PC1), species richness, and sampling effort. We assessed the sensitivity of each variable by comparing our observed effect estimates to the 95% quantiles of the corresponding null distributions of effect estimates from the simulated models ^98^. The observed effect of precipitation (PC1; *β* = 0.51) was greater than the 95% quantile of the null effect distribution (95% null *β* = 0.50), and the effect of temperature (PC2; *β* = 0.32) equaled the 92% quantile of the null effect distribution, indicating low sensitivity of our results to potential pseudo replication (Table S11).

### Complex display evolution predicts persistence following recent climate change

We examined how display complexity affected species persistence in changing climates by testing whether species that were observed within 2° latitude-wide hexagonal grid cells between 1970- 1999 were still observed within those grid cells between 2000-2024. We used GBIF occurrence records of the 12 *Schizocosa* species that had at least 30 occurrences between 1970-1999^60^. Species were considered to have complex displays if they had at least the median values of the number of courtship components and ornament exaggeration among all species used in our phylogenetic comparative analysis. This resulted in five species with complex displays and seven species with simple displays (see Table S16 for each species’ designation). For the 96 grid cells overlaying the historical occurrence records of the 12 species, we determined whether each species in the grid cell persisted into the present (Y/N; 189 persistence measurements). For each persistence measurement, we also calculated the change in temperature (the midpoint between the monthly *T*_min_ and *T*_max_^51^, averaged across all years in each time period) and change in_max_ precipitation (monthly precipitation^51^ averaged across all years in each time period) in the grid cell. The climate change calculations only included data from the three months in which the species is mature and reproductively active (based on collection dates in published literature and monthly frequencies of iNaturalist observations, Table S16). Species with simple displays experienced a much broader range of temperature changes in the last 50 years (-0.61 to 1.41°C) compared to species with complex displays (0.61 to 0.97°C). To standardize the degree of climate change between these groups, we limited our analysis to grid cells in which temperature increases were between 0.5 and 1.0°C, resulting in 75 grid cells and 153 persistence measurements among the 12 species. However, an analysis using all 96 grid cells showed similar results (Tables S13, S14). Following the approach of a recent study^32^, we used generalized linear mixed-effects models (‘*glmmTMB*’^61^) to test whether total display complexity (simple vs. complex) affected species’ probability of persistence in response to changing temperatures and precipitation. In all models, we included fixed effects of the grid cell’s change in temperature, change in precipitation, and their interactions with the display trait. We accounted for spatial variation in average persistence by including grid cell as a random effect. We also accounted for phylogenetic effects on average persistence by including random effects of species nested within the two major *Schizocosa* clades^34^. However, model comparison indicated that the best-supported models had only phylogenetic random effects (Table S12). We report results from these best supported models in the main text, but models with all random effects showed similar results (Table S13).

## Acknowledgments

N.T.L. was supported by a Saint Louis University Presidential Fellowship, a Saint Louis University Dissertation Fellowship, a University of Pittsburgh Ecology & Evolution Postdoctoral Fellowship, and a National Science Foundation Postdoctoral Research Fellowship in Biology (NSF DBI-2410506 to N.T.L.). J.P.W. was supported by a Saint Louis University Knoedler Research Grant, the University of South Florida, and a Genshaft Family Doctoral Fellowship. K.D.F.F. was supported by the Saint Louis University Research Institute. All authors were supported by NSF IOS-1656818 (to K.D.F.F.). E. Senthilkumaran helped with data collection. M.P. Moore, D.O. Elias, D. Gomez, L. Gath, E.A. Miller, K.D. Kohl, J.F. Stephenson, E. Hardison, J. Diaz, K. Peralta Martínez, and S.E. Nalley provided feedback on writing, analysis, and data visualization. We also thank all iNaturalist and GBIF contributors.

## Author Contributions

N.T.L. and K.D.F.F. designed the study. N.T.L. and J.P.W. performed the mating trial experiment. N.T.L. collected the interspecific and assemblage-level data, performed the statistical analyses, and wrote the manuscript with feedback from K.D.F.F. and J.P.W. Funding and project management was contribut

## References

1. West-Eberhard, M.J. (1983). Sexual selection, social competition, and speciation. The Quarterly Review of Biology 58, 155–183.

2. Mayr, E. (1963). Animal Species and Evolution (Harvard University Press).

3. Panhuis, T.M., Butlin, R., Zuk, M., and Tregenza, T. (2001). Sexual selection and speciation. Trends in Ecology and Evolution 16, 364–371. 10.1016/S0169-5347(01)02160-7.

4. Candolin, U. (2003). The use of multiple cues in mate choice. Biological Reviews 78, 575–595. 10.1017/S1464793103006158.

5. Hebets, E.A., Barron, A.B., Balakrishnan, C.N., Hauber, M.E., Mason, P.H., and Hoke, K.L. (2016). A systems approach to animal communication. Proceedings of the Royal Society B: Biological Sciences 283. 10.1098/rspb.2015.2889.

6. Hebets, E.A., and Papaj, D.R. (2005). Complex signal function: Developing a framework of testable hypotheses. Behavioral Ecology and Sociobiology 57, 197–214. 10.1007/s00265-004-0865-7.

7. Endler, J.A. (1992). Signals, signal conditions, and the direction of evolution. The American Naturalist 139, S125–S153.

8. Ryan, M.J., and Cummings, M.E. (2013). Perceptual Biases and Mate Choice. Annu. Rev. Ecol. Evol. Syst. 44, 437–459. 10.1146/annurev-ecolsys-110512-135901.

9. Ord, T.J., and Stamps, J.A. (2008). Alert signals enhance animal communication in “noisy” environments. Proceedings of the National Academy of Sciences 105, 18830–18835. 10.1073/pnas.0807657105.

10. Moss, J.B., Tumulty, J.P., and Fischer, E.K. (2023). Evolution of acoustic signals associated with cooperative parental behavior in a poison frog. Proceedings of the National Academy of Sciences 120, e2218956120. 10.1073/pnas.2218956120.

11. Bro-Jørgensen, J. (2010). Dynamics of multiple signalling systems: animal communication in a world in flux. Trends in Ecology and Evolution 25, 292–300. 10.1016/j.tree.2009.11.003.

12. Miles, M.C., Cheng, S., and Fuxjager, M.J. (2017). Biogeography predicts macro-evolutionary patterning of gestural display complexity in a passerine family. Evolution 71, 1406–1416. 10.1111/evo.13213.

13. Botero, C.A., Boogert, N.J., Vehrencamp, S.L., and Lovette, I.J. (2009). Climatic Patterns Predict the Elaboration of Song Displays in Mockingbirds. Current Biology 19, 1151–1155. 10.1016/j.cub.2009.04.061.

14. Broder, E.D., Elias, D.O., Rodríguez, R.L., Rosenthal, G.G., Seymoure, B.M., and Tinghitella, R.M. (2021). Evolutionary novelty in communication between the sexes. Biology Letters 17. 10.1098/rsbl.2020.0733.

15. West-Eberhard, M.J. (2005). Developmental plasticity and the origin of species differences. Proceedings of the National Academy of Sciences 102, 6543–6549. 10.1073/pnas.0501844102.

16. Moczek, A.P., Sultan, S., Foster, S., Ledón-Rettig, C., Dworkin, I., Nijhout, H.F., Abouheif, E., and Pfennig, D.W. (2011). The role of developmental plasticity in evolutionary innovation. Proceedings of the Royal Society B: Biological Sciences 278, 2705–2713. 10.1098/rspb.2011.0971.

17. Fisher, D.N., Kilgour, R.J., Siracusa, E.R., Foote, J.R., Hobson, E.A., Montiglio, P.O., Saltz, J.B., Wey, T.W., and Wice, E.W. (2021). Anticipated effects of abiotic environmental change on intraspecific social interactions. Biological Reviews 44. 10.1111/brv.12772.

18. Laiolo, P., Vögeli, M., and Tella, J.L. Song Diversity Predicts the Viability of Fragmented Bird Populations. PLoS ONE 3, e1822. 10.1371/%20journal.pone.0001822.

19. Tan, E.J., and Elgar, M.A. (2021). Motion: enhancing signals and concealing cues. Biology Open 10, bio058762. 10.1242/bio.058762.

20. Ligon, R.A., Diaz, C.D., Morano, J.L., Troscianko, J., Stevens, M., Moskeland, A., Laman, T.G., and Scholes, E. (2018). Evolution of correlated complexity in the radically different courtship signals of birds-of-paradise. PLOS Biology 16, e2006962. 10.1371/journal.pbio.2006962.

21. Hebets, E.A., Vink, C.J., Sullivan-Beckers, L., and Rosenthal, M.F. (2013). The dominance of seismic signaling and selection for signal complexity in Schizocosa multimodal courtship displays. Behavioral Ecology and Sociobiology 67, 1483–1498. 10.1007/s00265-013-1519-4.

22. Stafstrom, J.A., and Hebets, E.A. (2013). Female mate choice for multimodal courtship and the importance of the signaling background for selection on male ornamentation. Current Zoology 59, 200–209. 10.1093/czoolo/59.2.200.

23. Simpson, R.K., and McGraw, K.J. (2018). It’s not just what you have, but how you use it: solar- positional and behavioural effects on hummingbird colour appearance during courtship. Ecology Letters 21, 1413–1422. 10.1111/ele.13125.

24. White, T.E., Zeil, J., and Kemp, D.J. (2015). Signal design and courtship presentation coincide for highly biased delivery of an iridescent butterfly mating signal. Evolution 69, 14–25. 10.1111/evo.12551.

25. Bertossa, R.C. (2011). Morphology and behaviour: functional links in development and evolution. Philosophical Transactions of the Royal Society B: Biological Sciences 366, 2056–2068. 10.1098/rstb.2011.0035.

26. Leith, N.T., Macchiano, A., Moore, M.P., and Fowler-Finn, K.D. (2021). Temperature impacts all behavioral interactions during insect and arachnid reproduction. Current Opinion in Insect Science 45, 106–114. 10.1016/j.cois.2021.03.005.

27. García-Roa, R., Garcia-Gonzalez, F., Noble, D.W.A., and Carazo, P. (2020). Temperature as a modulator of sexual selection. Biological Reviews 95, 1607–1629. 10.1111/brv.12632.

28. Leith, N.T., Fowler-Finn, K.D., and Moore, M.P. (2022). Evolutionary interactions between thermal ecology and sexual selection. Ecology Letters 25, 1919–1936. 10.1111/ele.14072.

29. Rosenthal, M.F., and Elias, D.O. (2019). Nonlinear changes in selection on a mating display across a continuous thermal gradient. Proceedings of the Royal Society B: Biological Sciences 286, 20191450. 10.1098/rspb.2019.1450.

30. Partan, S.R. (2013). Ten unanswered questions in multimodal communication. Behavioral Ecology and Sociobiology 67, 1523–1539. 10.1007/s00265-013-1565-y.

31. Tuomainen, U., and Candolin, U. (2011). Behavioural responses to human-induced environmental change. Biological Reviews 86, 640–657. 10.1111/j.1469-185X.2010.00164.x.

32. Moore, M.P., Nalley, S.E., and Hamadah, D. (2024). An evolutionary innovation for mating facilitates ecological niche expansion and buffers species against climate change. Proceedings of the National Academy of Sciences 121, e2313371121. 10.1073/pnas.2313371121.

33. Servedio, M.R., and Boughman, J.W. (2017). The role of sexual selection in local adaptation and speciation. Annual Review of Ecology, Evolution, and Systematics 48, 85–109. 10.1146/annurev-ecolsys-110316-022905.

34. Starrett, J., McGinley, R.H., Hebets, E.A., and Bond, J.E. (2022). Phylogeny and secondary sexual trait evolution in *Schizocosa* wolf spiders (Araneae, Lycosidae) shows evidence for multiple gains and losses of ornamentation and species delimitation uncertainty. Molecular Phylogenetics and Evolution 169, 107397. 10.1016/j.ympev.2022.107397.

35. Stratton, G.E. (2005). Evolution of Ornamentation and Courtship Behavior in Schizocosa: Insights From a Phylogeny Based on Morphology (Araneae, Lycosidae). Journal of Arachnology 33, 347– 376. 10.1636/04-80.1.

36. Rosenthal, M.F., Wilkins, M.R., Shizuka, D., and Hebets, E.A. (2018). Dynamic changes in display architecture and function across environments revealed by a systems approach to animal communication*. Evolution 72, 1134–1145. 10.1111/evo.13448.

37. Davis, D.L. (1989). The Effect of Temperature on the Courtship Behavior of the Wolf Spider Schizocosa rovneri. American Midland Naturalist 122, 281–287.

38. Starrett, J., Bui, A., McGinley, R., Hebets, E.A., and Bond, J.E. (2021). Phylogenomic Variation at the Population-Species Interface and Assessment of Gigantism in a Model Wolf Spider Genus (Lycosidae, Schizocosa). Insect Systematics and Diversity 5, 5. 10.1093/isd/ixab016.

39. Hebets, E.A., Elias, D.O., Mason, A.C., Miller, G.L., and Stratton, G.E. (2008). Substrate-dependent signalling success in the wolf spider, Schizocosa retrorsa. Animal Behaviour 75, 605–615. 10.1016/j.anbehav.2007.06.021.

40. Stratton, G.E., and Lowrie, D.C. (1984). Courtship Behavior and Life Cycle of the Wolf Spider Schizocosa mccooki (Araneae , Lycosidae). Journal of Arachnology 12, 223–228.

41. McGinley, R.H., Starrett, J., Bond, J.E., and Hebets, E.A. (2023). Light Environment Interacts with Visual Displays in a Species-Specific Manner in Multimodal-Signaling Wolf Spiders. The American Naturalist 201, 472–490. 10.1086/722830.

42. Rundus, A.S., Sullivan-Beckers, L., Wilgers, D.J., and Hebets, E.A. (2011). Females are choosier in the dark: Environment-dependent reliance on courtship components and its impact on fitness. Evolution 65, 268–282. 10.1111/j.1558-5646.2010.01125.x.

43. Miller, G.L., Stratton, G.E., Miller, P.R., and Hebets, E.A. (1998). Geographical variation in male courtship behaviour and sexual isolation in wolf spiders of the genus Schizocosa. Animal Behaviour 56, 937–951. 10.1006/anbe.1998.0851.

44. Stratton, G.E., and Uetz, G.W. (1986). The inheritance of courtship behavior and its role as a reproductive isolating mechanism in two species of Schizocosa wolf spiders (Araneae; Lycosidae). Evolution 40, 129–141.

45. Fowler-Finn, K.D., Sullivan-Beckers, L., Runck, A.M., and Hebets, E.A. (2015). The complexities of female mate choice and male polymorphisms: Elucidating the role of genetics, age, and mate- choice copying. Current Zoology 61, 1015–1035. 10.1093/czoolo/61.6.1015.

46. Gordon, S.D., and Uetz, G.W. (2011). Multimodal communication of wolf spiders on different substrates: Evidence for behavioural plasticity. Animal Behaviour 81, 367–375. 10.1016/j.anbehav.2010.11.003.

47. Hebets, E.A., Bern, M., McGinley, R.H., Roberts, A., Kershenbaum, A., Starrett, J., and Bond, J.E. (2021). Sister species diverge in modality-specific courtship signal form and function. Ecology and Evolution 11, 852–871. 10.1002/ece3.7089.

48. Choi, N., Adams, M., Fowler-finn, K., Knowlton, E., Rosenthal, M., Rundus, A., Santer, R.D., Wilgers, D., Hebets, E.A., and Chester, W. (2022). Increased signal complexity is associated with increased mating success. Biology Letters 18, 20220052.

49. Choi, N., and Hebets, E.A. (2021). The effects of conspecific male density on the reproductive behavior of male Schizocosa retrorsa (Banks, 1911) wolf spiders (Araneae: Lycosidae). arac 49, 347–357. 10.1636/JoA-S-20-079.

50. Hebets, E.A. (2005). Attention-altering signal interactions in the multimodal courtship display of the wolf spider Schizocosa uetzi. Behavioral Ecology 16, 75–82. 10.1093/beheco/arh133.

51. Fick, S.E., and Hijmans, R.J. (2017). WorldClim 2: new 1-km spatial resolution climate surfaces for global land areas. International Journal of Climatology 37, 4302–4315. 10.1002/joc.5086.

52. Mittelbach, G.G., and Schemske, D.W. (2015). Ecological and evolutionary perspectives on community assembly. Trends in Ecology & Evolution 30, 241–247. 10.1016/j.tree.2015.02.008.

53. Parratt, S.R., Walsh, B.S., Metelmann, S., White, N., Manser, A., Bretman, A.J., Hoffmann, A.A., Snook, R.R., and Price, T.A.R. (2021). Temperatures that sterilize males better match global species distributions than lethal temperatures. Nature Climate Change 11, 481–484. 10.1038/s41558-021-01047-0.

54. Eliason, C.M., Nicolaï, M.P.J., Bom, C., Blom, E., D’Alba, L., and Shawkey, M.D. (2024). Transitions between colour mechanisms affect speciation dynamics and range distributions of birds. Nat Ecol Evol 8, 1723–1734. 10.1038/s41559-024-02487-5.

55. Maia, R., Rubenstein, D.R., and Shawkey, M.D. (2013). Key ornamental innovations facilitate diversification in an avian radiation. Proceedings of the National Academy of Sciences of the United States of America 110, 10687–10692. 10.1073/pnas.1220784110.

56. GBIF.org (2024). GBIF Occurrence Download. 10.15468/dl.jwztdv.

57. Chen, I.-C., Hill, J.K., Ohlemüller, R., Roy, D.B., and Thomas, C.D. (2011). Rapid Range Shifts of Species Associated with High Levels of Climate Warming. Science 333, 1024–1026. 10.1126/science.1206432.

58. van Heerwaarden, B., and Sgrò, C.M. (2021). Male fertility thermal limits predict vulnerability to climate warming. Nature Communications 12, 2214. 10.1038/s41467-021-22546-w.

59. Moore, M.P., Leith, N.T., Fowler-Finn, K.D., and Medley, K.A. (2024). Human-modified habitats imperil ornamented dragonflies less than their non-ornamented counterparts at local, regional, and continental scales. Ecology Letters 27, e14455. 10.1111/ele.14455.

60. GBIF.org (2025). GBIF Occurrence Download. 10.15468/dl.juu37m.

61. McGillycuddy, M., Popovic, G., Bolker, B.M., and Warton, D.I. (2025). Parsimoniously Fitting Large Multivariate Random Effects in glmmTMB. Journal of Statistical Software 112, 1–19. 10.18637/jss.v112.i01.

62. Weber, M.G., and Agrawal, A.A. (2012). Phylogeny, ecology, and the coupling of comparative and experimental approaches. Trends in Ecology and Evolution 27, 394–403. 10.1016/j.tree.2012.04.010.

63. Moore, M.P., Hersch, K., Sricharoen, C., Lee, S., Reice, C., Rice, P., Kronick, S., Medley, K.A., and Fowler-Finn, K.D. (2021). Sex-specific ornament evolution is a consistent feature of climatic adaptation across space and time in dragonflies. Proceedings of the National Academy of Sciences of the United States of America 118, 1–7. 10.1073/pnas.2101458118.

64. Pilakouta, N., and Ålund, M. (2021). Editorial: Sexual selection and environmental change: what do we know and what comes next? Current Zoology 67, 293–298. 10.1093/cz/zoab021.

65. Rozen-Rechels, D., Dupoué, A., Lourdais, O., Chamaillé-Jammes, S., Meylan, S., Clobert, J., and Le Galliard, J.F. (2019). When water interacts with temperature: Ecological and evolutionary implications of thermo-hydroregulation in terrestrial ectotherms. Ecology and Evolution 9, 10029– 10043. 10.1002/ece3.5440.

66. Sasson, D.A., Johnson, T.D., Scott, E.R., and Fowler-Finn, K.D. (2020). Short-term water deprivation has widespread effects on mating behaviour in a harvestman. Animal Behaviour 165, 97–106. 10.1016/j.anbehav.2020.04.026.

67. Ivy, T.M., Chadwick Johnson, J., and Sakaluk, S.K. (1999). Hydration benefits to courtship feeding in crickets. Proceedings of the Royal Society B: Biological Sciences 266, 1523–1527. 10.1098/rspb.1999.0810.

68. Beck, H.E., Zimmermann, N.E., McVicar, T.R., Vergopolan, N., Berg, A., and Wood, E.F. (2018). Present and future Köppen-Geiger climate classification maps at 1-km resolution. Sci Data 5, 180214. 10.1038/sdata.2018.214.

69. Leith, N.T., Miller, E.A., and Fowler-Finn, K.D. (2024). Thermoregulation enhances survival but not reproduction in a plant-feeding insect. Functional Ecology 38, 1344–1356. 10.1111/1365-2435.14546.

70. Endler, J.A., and Basolo, A.L. (1998). Sensory ecology, receiver biases and sexual selection. Trends in Ecology & Evolution 13, 415–420. 10.2307/1446782.

71. Cummings, M.E. (2007). Sensory trade-offs predict signal divergence in surfperch. Evolution 61, 530–545. 10.1111/j.1558-5646.2007.00047.x.

72. Leith, N.T., and Moore, M.P. (2024). Heat-absorbing sexual coloration co-adapts with increased heat tolerance in dragonflies. Front. Ethol. 3, 1447637. 10.3389/fetho.2024.1447637.

73. Fuxjager, M.J., Ryder, T.B., Moody, N.M., Alfonso, C., Balakrishnan, C.N., Barske, J., Bosholn, M., Boyle, W.A., Braun, E.L., Chiver, I., et al. (2023). Systems biology as a framework to understand the physiological and endocrine bases of behavior and its evolution—From concepts to a case study in birds. Hormones and Behavior 151, 105340. 10.1016/j.yhbeh.2023.105340.

74. Levis, N.A., Isdaner, A.J., and Pfennig, D.W. (2018). Morphological novelty emerges from pre- existing phenotypic plasticity. Nat Ecol Evol 2, 1289–1297. 10.1038/s41559-018-0601-8.

75. Endler, J.A., Butlin, R.K., Guilford, T., and Krebs, J.R. (1993). Some general comments on the evolution and design of animal communication systems. Philosophical Transactions of the Royal Society of London. Series B: Biological Sciences 340, 215–225. 10.1098/rstb.1993.0060.

76. Herrmann, S., and Roberts, J. (2017). Dehydration resistance and tolerance in the brush-legged wolf spider (Schizocosa ocreata): A comparison of survivorship, critical body water content, and water loss rates between sexes. Canadian Journal of Zoology 95, 417–423.

77. Chabaud, C., Brusch, G.A., Pellerin, A., Lourdais, O., and Le Galliard, J.-F. (2023). Prey consumption does not restore hydration state but mitigates the energetic costs of water deprivation in an insectivorous lizard. Journal of Experimental Biology 226. 10.1242/jeb.246129.

78. Nychka, D., Furrer, R., Paige, J., and Sain, S. (2021). fields: Tools for spatial data. Version 14.1 (University Corporation for Atmospheric Research).

79. R Core Team (2025). R: A language and environment for statistical computing. Version 4.5.1 (R Foundation for Statistical Computing).

80. Uetz, G.W., and Norton, S. (2007). Preference for male traits in female wolf spiders varies with the choice of available males, female age and reproductive state. Behavioral Ecology and Sociobiology 61, 631–641. 10.1007/s00265-006-0293-y.

81. Uetz, G.W., Stoffer, B., Lallo, M.M., and Clark, D.L. (2017). Complex signals and comparative mate assessment in wolf spiders: results from multimodal playback studies. Animal Behaviour 134, 283– 299. 10.1016/j.anbehav.2017.02.007.

82. Gibson, J.S., and Uetz, G.W. (2008). Seismic communication and mate choice in wolf spiders: components of male seismic signals and mating success. Animal Behaviour 75, 1253–1262. 10.1016/j.anbehav.2007.09.026.

83. Schneider, C.A., Rasband, W.S., and Eliceiri, K.W. (2012). NIH Image to ImageJ: 25 years of image analysis. Nat Methods 9, 671–675. 10.1038/nmeth.2089.

84. Wilkins, M.R., Shizuka, D., Joseph, M.B., Hubbard, J.K., and Safran, R.J. (2015). Multimodal signalling in the North American barn swallow: a phenotype network approach. Proceedings of the Royal Society B: Biological Sciences 282, 20151574. 10.1098/rspb.2015.1574.

85. Epskamp, S., Cramer, A.O.J., Waldorp, L.J., Schmittmann, V.D., and Borsboom, D. (2012). qgraph: Network Visualizations of Relationships in Psychometric Data. Journal of Statistical Software 48, 1– 18. 10.18637/jss.v048.i04.

86. Farine, D.R. (2014). Measuring phenotypic assortment in animal social networks: weighted associations are more robust than binary edges. Animal Behaviour 89, 141–153. 10.1016/j.anbehav.2014.01.001.

87. Revelle, W. (2023). psych: Procedures for Personality and Psychological Research. Version 2.3.9.

88. Schielzeth, H. (2010). Simple means to improve the interpretability of regression coefficients. Methods in Ecology and Evolution 1, 103–113. 10.1111/j.2041-210x.2010.00012.x.

89. Revell, L.J. (2012). phytools: An R package for phylogenetic comparative biology (and other things). Methods in Ecology and Evolution 3, 217–223. 10.1111/j.2041-210X.2011.00169.x.

90. Tung Ho, L.S., and Ané, C. (2014). A linear-time algorithm for gaussian and non-gaussian trait evolution models. Systematic Biology 63, 397–408. 10.1093/sysbio/syu005.

91. van der Bijl, W. (2018). phylopath: Easy phylogenetic path analysis in R. PeerJ 6, e4718. 10.7717/peerj.4718.

92. Lajeunesse, A., and Fourcade, Y. (2023). Temporal analysis of GBIF data reveals the restructuring of communities following climate change. Journal of Animal Ecology 92, 391–402. 10.1111/1365-2656.13854.

93. Gaüzère, P., Jiguet, F., and Devictor, V. (2015). Rapid adjustment of bird community compositions to local climatic variations and its functional consequences. Global Change Biology 21, 3367–3378. 10.1111/gcb.12917.

94. Hijmans, R.J., Barbosa, M., Bivand, R., Brown, A., Chirico, M., Cordano, E., Dyba, K., Pebesma, E., Rowlingson, B., and Sumner, M.D. (2025). terra: Spatial Data Analysis. Version 1.8–54.

95. Pebesma, E. (2018). Simple Features for R: Standardized Support for Spatial Vector Data. The R Journal 10, 439–446.

96. Pinherio, J., Bates, D., DebRoy, S., Sarkar, D., and Team, R.C. (2020). nlme: Linear and Nonlinear mixed effects models. R package version 3.1–151.

97. Hawkins, B.A., Leroy, B., Rodríguez, M.Á., Singer, A., Vilela, B., Villalobos, F., Wang, X., and Zelený, D. (2017). Structural bias in aggregated species-level variables driven by repeated species co- occurrences: a pervasive problem in community and assemblage data. Journal of Biogeography 44, 1199–1211. 10.1111/jbi.12953.

98. Friedman, N.R., and Remeš, V. (2017). Ecogeographical gradients in plumage coloration among Australasian songbird clades. Global Ecology and Biogeography 26, 261–274. 10.1111/geb.12522.

99. DeVito, J., and Formanowicz, D.R. (2003). The effects of size, sex, and reproductive condition on thermal and desiccation stress in a riparian spider (Pirata sedentarius, Araneae, Lycosidae). Journal of Arachnology 31, 278–284. 10.1636/02-20.

100. Revell, L.J. (2010). Phylogenetic signal and linear regression on species data. Methods in Ecology and Evolution 1, 319–329. 10.1111/j.2041-210x.2010.00044.x.

