## Supplemental Information for "Mating display plasticity predicts the biogeography of complex communication and population persistence in changing climates"

**Author Contributions:** N.T.L. and K.D.F.F. designed the study. N.T.L. and J.P.W. performed the mating trial experiment. N.T.L. collected the interspecific and assemblage-level data, performed the statistical analyses, and wrote the manuscript with feedback from K.D.F.F. and J.P.W. Funding and project management was contributed by K.D.F.F.

**Competing Interest Statement:** The authors declare no competing interests.

**Keywords:** sexual selection; mate choice; phenotypic plasticity; sensory ecology; climate change

### Supplemental Information

### Supplemental Methods

**Comparing laboratory treatments to environmental conditions in the field.** To assess the ecological relevance of our water availability treatments, we compared body water content between treatments to variation in body water content among adult spiders at our field site near St. Louis, Missouri, USA (Lat: 38.554291; Long: -90.369585; Figure S1A). On eight days between May 10 and May 26, 2021, we used pre-weighed glass vials to collect male and female spiders at four sample locations in the field site. We sampled from two locations on hilltops and two locations near streambeds because fine-scale differences in local hydrology can influence wolf spider hydration[99]. All sampling locations were separated by at least 100 meters. After collection, we immediately froze spiders in the laboratory at -20ºC for at least 24 hours. We then weighed all individuals before and after drying at 60°C for 24 hours[99] to calculate body water content.

We used the same range of trial temperatures that were previously used to test how temperature affects courtship behavior and mating success in a closely related *Schizocosa* species [29]. The year after the laboratory experiment described in our study (2022), we also deployed three iButtons (DS1923-F5# Hygrochon temperature and humidity loggers) at our field site to verify that the trial temperatures used in our study were within the range of microclimates that *S. ocreata* regularly experience in the field (Figure S1B). The three iButtons were placed approximated 150 meters apart, rested on top of the leaf litter (where wolf spiders engage in courtship behavior[45,46]), and were secured in place with a metal stake placed in the soil. We programmed the iButtons to measure microclimate temperatures every hour during peak mating season (May 2 to July 8, 2022).

**Exploring nonlinear effects of temperature on mating behavior.** We used thin-plate splines (‘*fields*’[78]) to examine nonlinear relationships between sexual traits and mating success across continuous variation in temperatures. We analyzed data separately for each water treatment. For each of the 12 male sexual traits (Figure 2A; Figure S2), we constructed a 2D surface plot with temperature as the x-axis, the sexual trait variable as the y-axis, and mating success (yes/no) as the z-axis. Visualizing the 2D surface plots revealed that increased temperatures generated nonlinear changes in selection on the number of tapping bouts, percussion period, ornamental tuft area, femur darkness, and body size (Figures S3 and S4). However, the effect of each trait on mating success always increased or decreased monotonically with increased temperature (Figures S3 and S4). Furthermore, when nonlinear changes in selection occurred (indicated by curved contour lines on the surface plot), we observed an inflection point in the trait’s effect on mating success near the median temperature value used to delineate our two main temperature treatments (25.3ºC). Taken together, these observations suggest that sorting our trials into two temperature treatments (cold: less than or equal to 25.3ºC; hot: greater than 25.3ºC) adequately captured changes in the function of mating displays across temperatures.

**Characterizing variation in sexual traits among taxa.** We used previously published courtship descriptions and videos to quantify the total number of male courtship behaviors used by each taxon in the phylogeny[21,35–47,49,50] (Table S15). We only used courtship descriptions that were published in peer-reviewed articles and included information on gestural behaviors, vibrational acoustic behaviors, and geographic location. For each species represented in the phylogeny, we compiled all courtship descriptions that met the above criteria. We then assigned each taxon in the phylogeny the courtship description from the article on its species that was closest to the taxon's geographic coordinates. From each courtship description, we determined the number of unique behaviors incorporated into the courtship repertoire. Importantly, our hypothesis for how climate affects courtship evolution involves constraints on the behaviors that produce courtship stimuli, not the transmission or perception of those stimuli. A single behavior may produce only a visual stimulus (e.g., leg waving), only a vibrational stimulus (e.g., palpal stridulation), or both visual and vibrational stimuli (e.g., percussive strikes/body bounces). We quantified courtship repertoires by counting the number of behaviors that produce at least one stimulus, regardless of modality.

To quantify the degree of ornamentation for each taxon, we used the standardized ornamentation score computed by Starrett and colleagues[34]. This score adds together the proportion of the tibia and metatarsus covered by ornamental bristles, the density of bristles in the ornament, and the proportion of forelimb area covered in dark pigmentation. Prior work suggests that these traits have redundant effects for eliciting female responses, so the cumulative degree of ornamentation among all traits is likely more important for ornament function than the expression of different trait combinations within the ornament[34].

**Phylogenetic generalized least squares with total display complexity as a continuous trait.** We tested whether taxa have evolved increased display complexity (varying continuously across taxa) using a phylogenetic generalized least squares model (PGLS; '*phytools*'[89]). We included the total display complexity score as response variables and temperature (pPC1), precipitation (pPC2), and their interaction as the explanatory variables. The phylogenetic generalized linear modeling framework we used does not compute confidence intervals for Poisson distributed responses[90]; we therefore $\log_e$-transformed(+1) the total complexity score to improve model fit. Total complexity was then z-transformed to standardize regression coefficients[88]. To assess which model of evolutionary history we should use to account for any phylogenetic non-independence in our PGLS analyses[100], we used AICc to compare among a Brownian motion model, models using Pagel's $\lambda$ branch-length transformation, and a model without phylogeny. However, models with Pagel's $\lambda$ did not converge. We performed the PGLS with a star phylogeny, as this model showed the greatest support (Table S6).

**Model comparisons for phylogenetic path analysis.** We performed a phylogenetic path analysis under Brownian motion ('*phylopath*'[91]). We used Brownian motion because *phylopath* requires a specified model of evolution[91] and models with Pagel's $\lambda$ branch-length transformation did not converge in our PGLS analysis. We constructed six hypothesized path models that differed in the inclusion of direct and indirect effects of climate on the evolution of behavioral courtship complexity and ornament exaggeration. We created models where temperature and precipitation directly affected either behavioral complexity (Figure S8B) or ornament exaggeration (Figure S8A); models where temperature and precipitation only indirectly affected behavioral complexity (Figure S8C) or ornament exaggeration (Figure S8D); and models with a combination of direct and indirect effects of climate variables on each trait type (Figure S8E and S8F). We performed one path analyses with behavioral complexity and ornament exaggeration as binary variables (Figure S8) and one path analyses with each trait as a continuous variable (Figure S9; see *Characterizing variation in sexual traits among taxa*). For the analysis with continuous traits, we $\log_e$-transformed(+1) the number of courtship behaviors and ornament exaggeration score and then z-transformed each variable to standardize path coefficients[88]. If two or more models received equal support (ΔCICc < 2), we performed model averaging to determine path coefficients and 95% confidence intervals.

**Community-level spatial analysis with non-overlapping assemblages.** We generated a presence/absence matrix of 2° latitude-wide hexagonal grid cells for each species across the geographic extents of the occurrence records ('*terra*'[94]). For each cell, we determined the mean display complexity score among all species observed in the cell. For species that were represented by multiple taxa in the phylogeny[34], we used the mean display complexity score among every taxon associated with that species. To accurately characterize assemblage-level mean display complexity, we only included cells with at least two species[92], resulting in 142 assemblages. We also quantified species richness and the total number of observations included in each cell. Following the justification outlined in our phylogenetic comparative analysis, we determined the mean values of four bioclimatic variables across the entire area of each cell[51]: (i) mean temperature of the wettest quarter; (ii) maximum temperature of the warmest month; (iii) precipitation of the wettest quarter; and (iv) precipitation of the driest month. Both precipitation variables were $\log_e$-transformed before analysis. A principal components analysis on these bioclimatic variables yielded two components that collectively explained 77% of the variation in the dataset. We varimax-rotated these components to improve the interpretability of our results[87], resulting in one principal component that showed a strong positive association with temperature (PC1) and another principal component that showed a strong positive association with precipitation (PC2; Table S9).

We used spatially explicit generalized least squares models (spatial GLS; '*nlme*'[96]) to assess the relative importance of temperature, precipitation, and species richness for shaping variation in multi-component displays among assemblages. We fit the $\log_e$-transformed(+1) assemblage-mean display complexity scores as the response, with effects of temperature (PC1), precipitation (PC2), the interaction between PC1 and PC2, and $\log_e$-transformed species richness. To account for variation in sampling effort, we included the $\log_e$-transformed total number of observations in the grid cell as covariate. All variables were then z-transformed to standardize regression coefficients[88]. We used rational quadratic spatial correlation structure because it received better support than models with alternative correlation structures (determined with ΔAICc; Table S10). Results for this spatial analysis with non-overlapping assemblages are reported in Table S11.

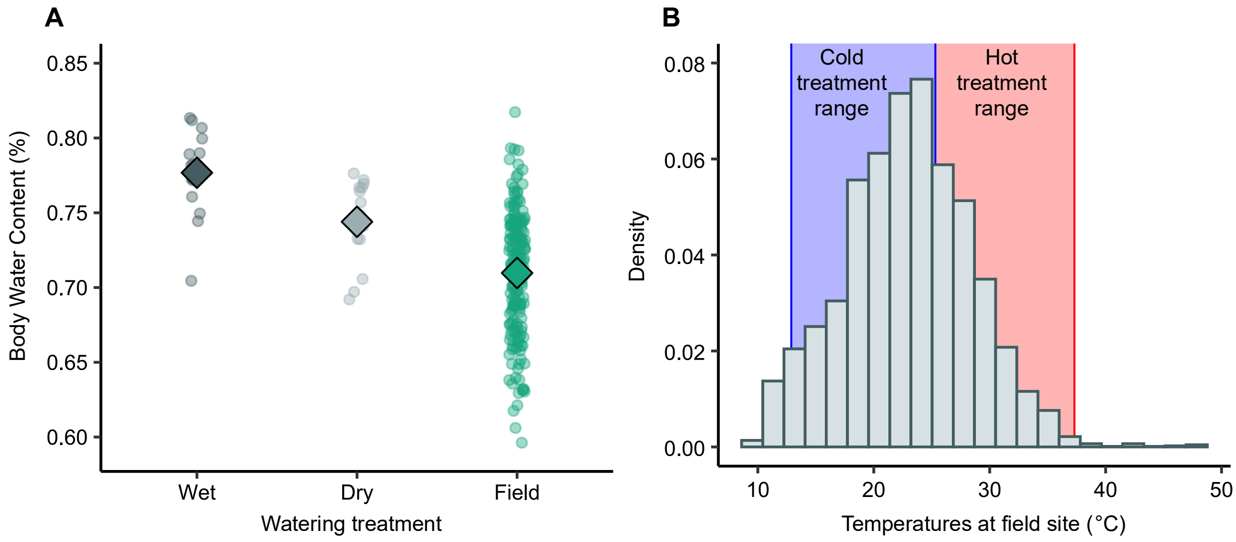


**Figure S1. Comparisons between laboratory environmental manipulations and environmental variation in the field during peak mating activity. (A)** Individuals in the wet treatment (dark grey points) had higher body water content than individuals in the dry treatment (light grey points). However, body water content measured in the field (green points) was much more variable, and all spiders in the lab maintained high body water content compared to those in the field. Diamonds represent median values. **(B)** Histogram of temperatures measured by iButton temperature loggers during the mating season (May 2-July 8) at the field site where *S. ocreata* were collected as juveniles.

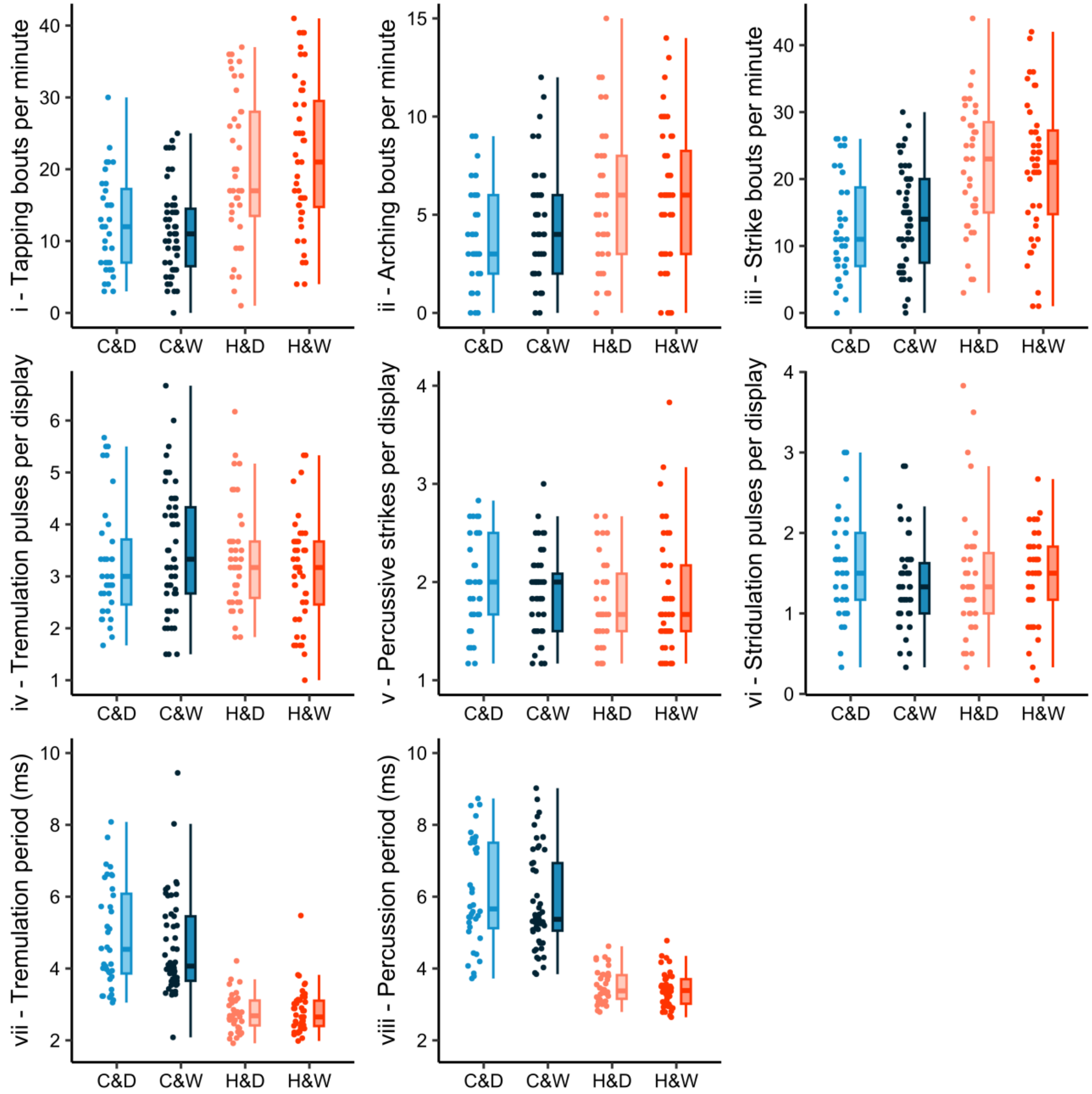


**Figure S2. Variation in male sexual traits in the cold-and-dry, cold-and-wet, hot-and-dry, and hot-and-wet treatments.** Points indicate trait values from a single male in a single trial.

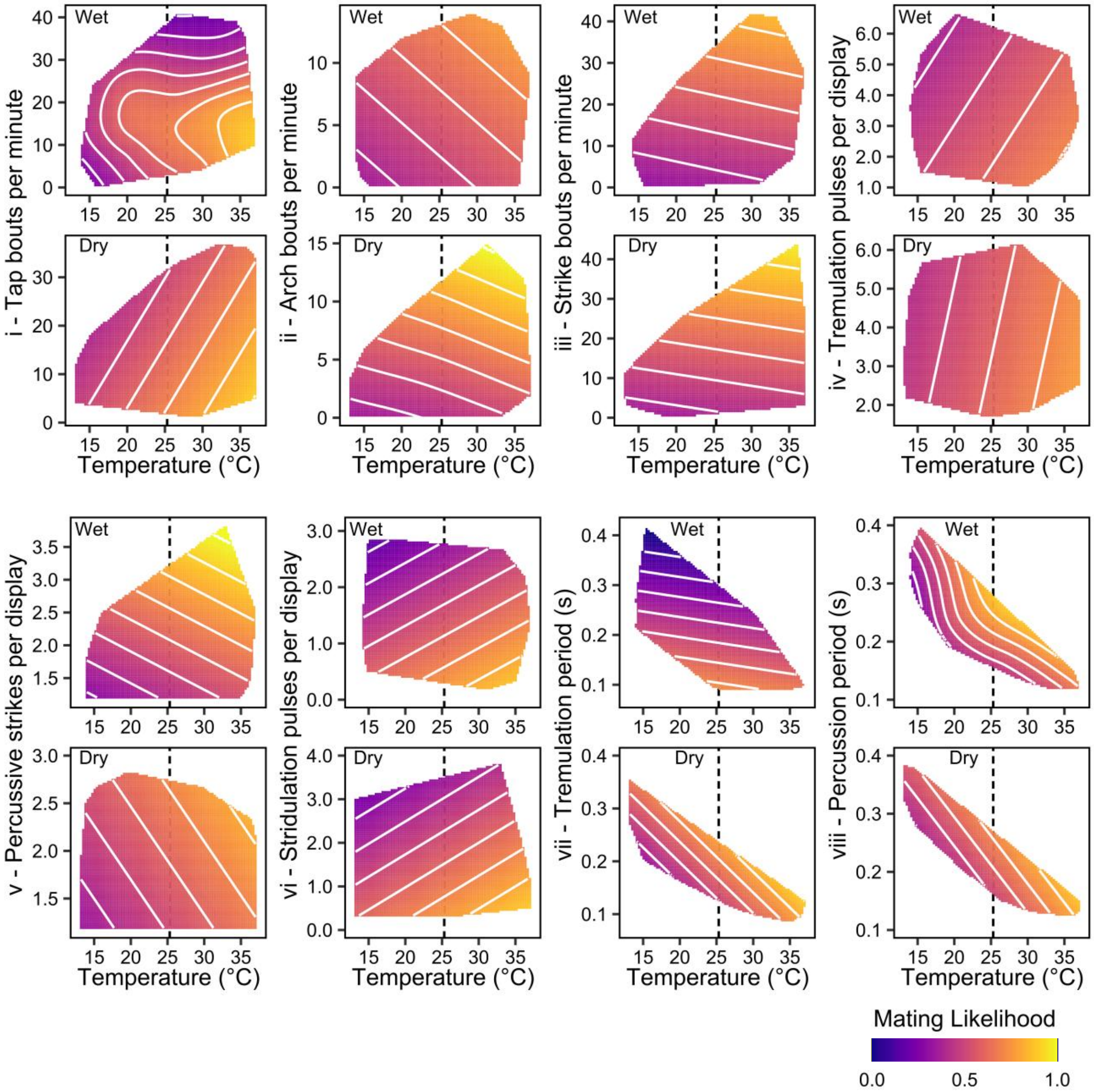


**Figure S3. Nonlinear effects of temperature on the function of courtship traits for mating success.** The 2D surfaces were constructed using thin-plate splines. Curved contour lines indicate nonlinear effects of temperature on the relationship between sexual traits and mating success. The dashed line indicates the median trial temperature. Temperatures to the left of or equal to the dashed line were included in the cold-and-dry and cold-and-wet treatments, while values to the right of the dashed line were included in the hot-and-dry and hot-and-wet treatments.

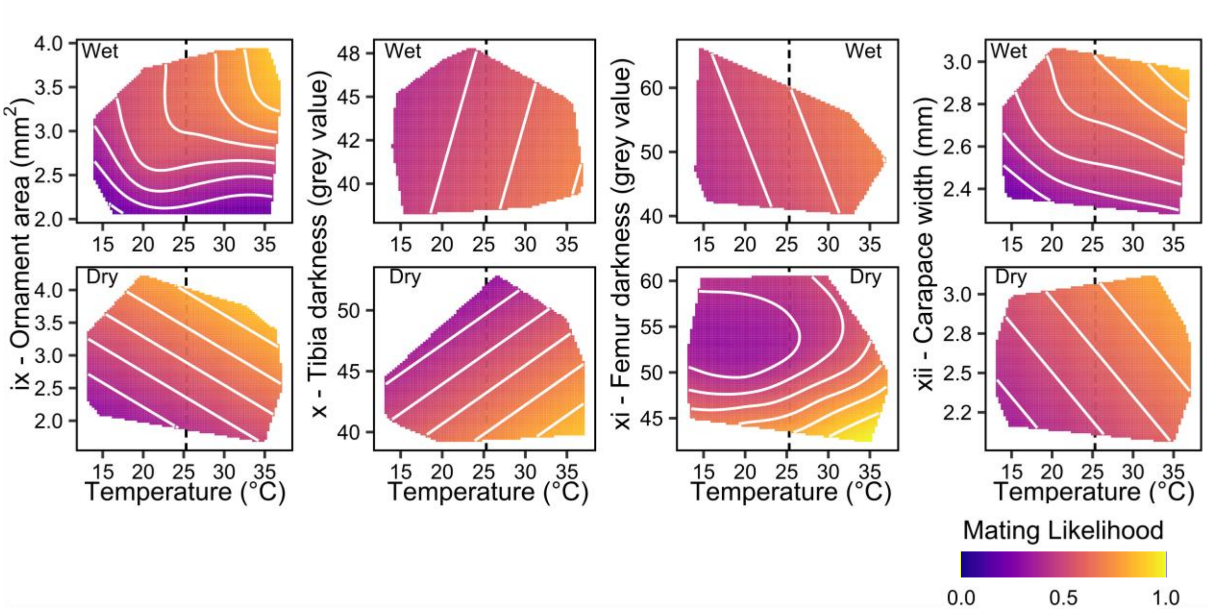


**Figure S4. Nonlinear effects of temperature on the function of morphological sexual traits for mating success.** The 2D surfaces were constructed using thin-plate splines. Curved contour lines indicate nonlinear effects of temperature on the relationship between sexual traits and mating success. The dashed line indicates the median trial temperature. Temperatures to the left of or equal to the dashed line were included in the cold-and-dry and cold-and-wet treatments, while values to the right of the dashed line were included in the hot-and-dry and hot-and-wet treatments.

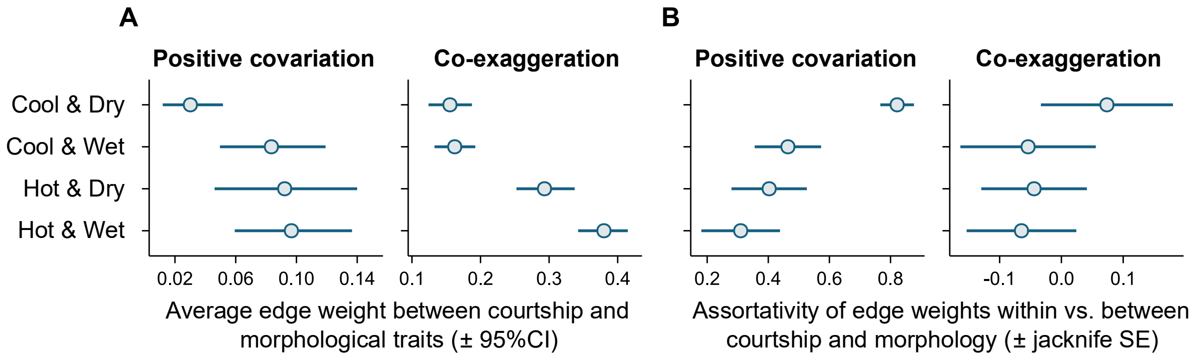


**Figure S5.** Indices of co-exaggeration and positive covariation between behavioral and morphological traits. Larger values of between-type edge weight and smaller values of assortativity indicate greater covariation between behavioral and morphological traits. Points are network-level indices. For **(A)**, bars are 95% CIs from 1000 bootstrapped iterations. For **(B)**, bars are jacknife SEs.

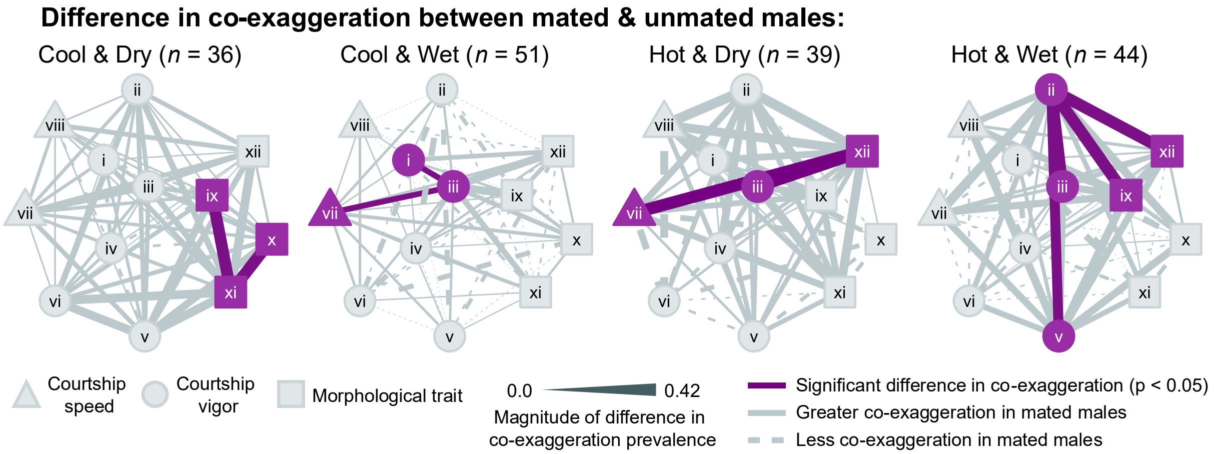


**Figure S6.** Phenotype networks illustrating the effects of sexual trait co-exaggeration on mating success in each environmental treatment. Courtship speed traits are shown in triangles, courtship vigor traits are shown in circles, and morphological sexual traits are in squares. Thicker edges between traits indicate greater differences in co-exaggeration prevalence between mated and unmated males. Solid edges indicate trait pairs with greater co-exaggeration in mated males, and dashed edges are trait pairs with less co-exaggeration in mated males. Purple edges indicate trait pairs with significant differences in co-exaggeration between mated and unmated males.

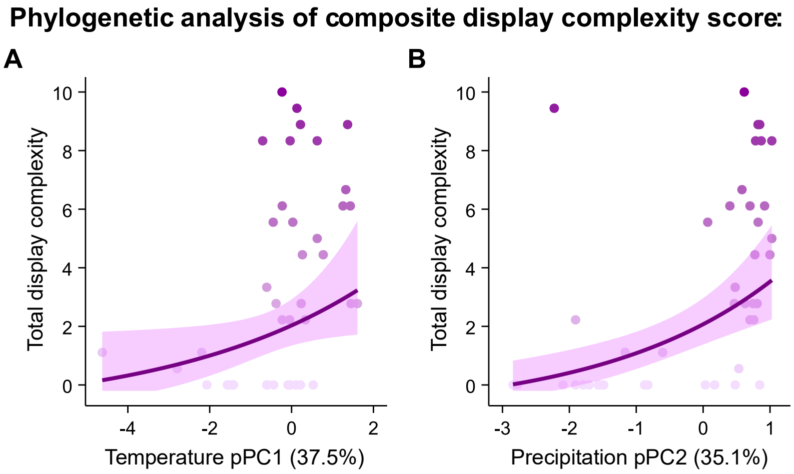


**Figure S7. Species in hotter and wetter climates evolve increased display complexity.** Total display complexity increases in taxa that have evolved in hotter **(A)** and wetter **(B)** climates. Points indicate individual taxa (*n* = 42), bolded lines are model estimates, and bands are 95% confidence intervals from a phylogenetic generalized least squares model.

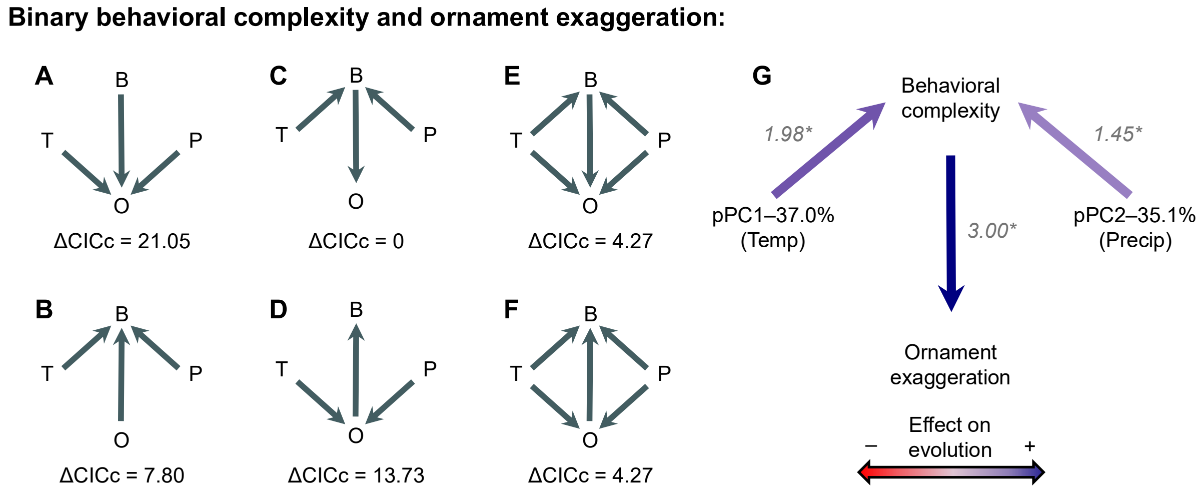


**Figure S8. Direct and indirect effects of climate on the coevolution of behavioral complexity and ornament exaggeration (behavior and ornamentation treated as binary variables). (A-F)** Path diagrams showing hypothesized causal evolutionary relationships among the number of additional courtship behaviors ("B"), ornament exaggeration ("O"), temperature ("T"), and precipitation ("P"). **(G)** Path diagram illustrating the standardized effect estimates and 95% confidence intervals for the best supported phylogenetic path model **(C)**. Blue path arrows reflect positive relationships between traits, while red arrows reflect negative effects. Arrows with more saturated coloration indicate stronger effect sizes. Significant parameter estimates indicated with asterisks.

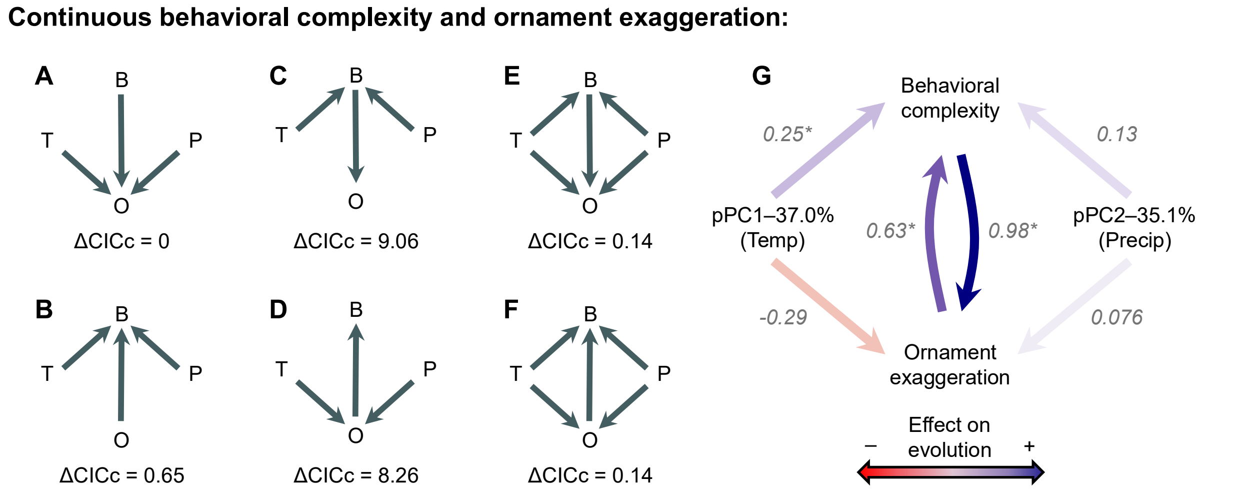


**Figure S9. Direct and indirect effects of climate on the coevolution of behavioral complexity and ornament exaggeration (behavior and ornamentation treated as continuous variables). (A-F)** Path diagrams showing hypothesized causal evolutionary relationships among the number of additional courtship behaviors ("B"), ornament exaggeration ("O"), temperature ("T"), and precipitation ("P"). **(G)** Path diagram illustrating the model-averaged standardized effect estimates and 95% confidence intervals for the four best supported phylogenetic path models **(A, B, E, F)**. Blue path arrows reflect positive relationships between traits, while red arrows reflect negative effects. Arrows with more saturated coloration indicate stronger effect sizes. Significant parameter estimates indicated with asterisks.

**Table S1.** Indices of co-exaggeration and positive covariation between behavioral and morphological traits. Larger values of between-type edge weight and smaller values of assortativity indicate greater covariation between behavioral and morphological traits.

| Network type | Environemntal treatment | Average between-type edge weight (95% CI) | Assortativity within trait types, $r_d$ (Jackknife SE) |
|---|---|---|---|
| Co-exaggeration | Cold & Dry | 0.16 (0.12 – 0.19) | 0.07 (-0.03 – 0.18) |
| | Cold & Wet | 0.16 (0.13 – 0.19) | -0.05 (-0.16 – 0.06) |
| | Hot & Dry | 0.29 (0.25 – 0.34) | -0.04 (-0.13 – 0.04) |
| | Hot & Wet | 0.38 (0.34 – 0.41) | -0.06 (-0.15 – 0.02) |
| Positive covariation | Cold & Dry | 0.03 (0.01 – 0.05) | 0.82 (0.77 – 0.88) |
| | Cold & Wet | 0.08 (0.05 – 0.12) | 0.46 (0.36 – 0.57) |
| | Hot & Dry | 0.09 (0.05 – 0.14) | 0.40 (0.28 – 0.53) |
| | Hot & Wet | 0.10 (0.06 – 0.14) | 0.31 (0.18 – 0.44) |

**Table S2.** Principal component loading factors for male sexual traits in each environmental treatment. The principal components analyses were performed on the correlation matrices of the 12 sexual traits in each treatment. The percentage of variance explained by each principal component is indicated in parentheses. Trait numerals correspond to numerals in Figure 2A. Shaded PCs indicate components that had the strongest effect on mating success in each treatment. In the Hot & Wet treatment, two PCs are shaded because both components had significant effects on mating success. Shaded loadings indicate traits that load strongly (magnitude > 0.4) and are favored by estimated selection on the principal components that best predicted mating success (negative loadings indicate favored traits for principal components that had negative effects on mating success).

| | Cool & Dry | | | | | Cool & Wet | | | | |
|---|---|---|---|---|---|---|---|---|---|---|
| Trait | PC1 (29%) | PC2 (24%) | PC3 (11%) | PC4 (10%) | PC5 (8%) | PC1 (27%) | PC2 (18%) | PC3 (13%) | PC4 (10%) | PC5 (8%) |
| i - TB | 0.63 | 0.13 | -0.48 | 0.33 | 0.25 | 0.53 | -0.28 | -0.40 | -0.38 | -0.08 |
| ii - AB | 0.68 | 0.00 | 0.40 | -0.23 | 0.34 | 0.45 | -0.15 | 0.33 | 0.66 | 0.02 |
| iii - SB | 0.80 | 0.30 | -0.09 | -0.33 | -0.02 | 0.83 | -0.10 | 0.15 | -0.16 | 0.04 |
| iv - Tp | -0.18 | -0.22 | 0.59 | 0.32 | 0.64 | 0.00 | -0.20 | -0.42 | 0.60 | 0.50 |
| v - Ps | 0.47 | 0.12 | 0.52 | -0.52 | -0.17 | 0.20 | -0.06 | 0.52 | -0.32 | 0.63 |
| vi - Sp | 0.12 | -0.26 | 0.52 | 0.48 | -0.55 | -0.02 | -0.06 | 0.69 | 0.15 | -0.41 |
| vii - Tr | 0.75 | 0.44 | -0.02 | 0.35 | -0.02 | 0.81 | -0.34 | 0.01 | -0.01 | 0.04 |
| viii - Pr | 0.80 | 0.37 | 0.05 | 0.34 | -0.06 | 0.82 | -0.38 | 0.10 | 0.01 | -0.08 |
| ix - OR | -0.38 | 0.79 | 0.04 | -0.01 | -0.01 | 0.60 | 0.56 | -0.23 | 0.20 | -0.17 |
| x - TD | -0.56 | 0.68 | 0.06 | 0.07 | 0.16 | 0.11 | 0.70 | 0.43 | 0.08 | 0.09 |
| xi - FD | -0.28 | 0.66 | 0.22 | 0.25 | -0.11 | 0.22 | 0.71 | 0.00 | -0.14 | 0.29 |
| xii - BS | -0.16 | 0.89 | 0.07 | -0.18 | -0.01 | 0.51 | 0.64 | -0.30 | 0.03 | -0.15 |
| | Hot & Dry | | | | | Hot & Wet | | | | |
| Trait | PC1 (28%) | PC2 (16%) | PC3 (15%) | PC4 (11%) | PC5 (8%) | PC1 (25%) | PC2 (16%) | PC3 (12%) | PC4 (11%) | PC5 (10%) |
| i - TB | 0.25 | 0.40 | -0.70 | 0.05 | -0.15 | -0.25 | 0.41 | 0.67 | 0.20 | 0.34 |
| ii - AB | 0.37 | 0.11 | 0.47 | -0.54 | -0.22 | 0.53 | -0.16 | -0.12 | -0.42 | 0.21 |
| iii - SB | 0.69 | 0.41 | -0.13 | -0.37 | -0.11 | 0.36 | 0.66 | 0.36 | -0.20 | 0.11 |
| iv - Tp | -0.23 | -0.17 | 0.45 | 0.56 | 0.26 | -0.18 | -0.60 | 0.13 | 0.21 | 0.54 |
| v - Ps | -0.17 | 0.06 | 0.39 | -0.59 | 0.57 | 0.53 | 0.43 | -0.21 | -0.17 | -0.40 |
| vi - Sp | -0.48 | 0.37 | 0.50 | 0.14 | -0.36 | 0.39 | 0.03 | 0.65 | -0.22 | 0.04 |
| vii - Tr | 0.09 | 0.78 | 0.04 | 0.21 | 0.41 | -0.07 | 0.52 | -0.49 | 0.18 | 0.47 |
| viii - Pr | 0.15 | 0.83 | 0.13 | 0.26 | 0.05 | -0.32 | 0.65 | -0.28 | 0.15 | 0.22 |
| ix - OR | 0.79 | -0.08 | 0.37 | 0.18 | -0.15 | 0.78 | 0.00 | -0.08 | -0.17 | 0.34 |
| x - TD | 0.82 | -0.11 | 0.09 | 0.00 | 0.23 | 0.65 | -0.14 | -0.05 | 0.65 | 0.08 |
| xi - FD | 0.74 | -0.34 | -0.27 | 0.22 | 0.28 | 0.49 | 0.11 | 0.16 | 0.68 | -0.35 |
| xii - BS | 0.73 | -0.09 | 0.42 | 0.24 | -0.20 | 0.81 | -0.14 | -0.16 | -0.01 | 0.23 |

**Table S3.** Effects of principal components (representing covariation in sexual traits) on mating success in each environmental treatment. Exact mating temperatures varied continuously within each categorical "cold" (12.9-25.3ºC) and "hot" (25.4-37.3ºC) treatment; exact temperature was therefore included as a covariate in all models. Results were obtained from binomial generalized linear models. Significance was assessed using likelihood ratio tests comparing a full model to a model in which the variable of interest was removed. Standardized parameter estimates ($\beta$) and confidence intervals were obtained from the full models.

| Treatment | Variable | LR $\chi_1^2$ | df | P | $\beta$ | Low 95% CI | High 95% CI |
|---|---|---|---|---|---|---|---|
| Cool & Dry | PC1 | 0.03 | 1 | 0.8671 | -0.16 | -2.11 | 1.76 |
| | PC2 | 0.45 | 1 | 0.5009 | 0.34 | -0.65 | 1.43 |
| | PC3 | 0.18 | 1 | 0.6698 | 0.15 | -0.56 | 0.88 |
| | PC4 | 1.07 | 1 | 0.3011 | -0.49 | -1.53 | 0.43 |
| | PC5 | 0.59 | 1 | 0.4413 | -0.30 | -1.11 | 0.47 |
| | Exact temperature | 0.04 | 1 | 0.8375 | 0.22 | -1.94 | 2.42 |
| Cool & Wet | PC1 | 1.77 | 1 | 0.1830 | 0.77 | -0.36 | 2.03 |
| | PC2 | 0.50 | 1 | 0.4812 | -0.25 | -1.01 | 0.44 |
| | PC3 | 2.04 | 1 | 0.1528 | -0.46 | -1.17 | 0.17 |
| | PC4 | 3.14 | 1 | 0.0762 | -0.56 | -1.27 | 0.06 |
| | PC5 | 0.35 | 1 | 0.5519 | 0.19 | -0.44 | 0.86 |
| | Exact temperature | 0.03 | 1 | 0.8681 | -0.10 | -1.32 | 1.10 |
| Hot & Dry | PC1 | 0.96 | 1 | 0.3262 | 0.42 | -0.42 | 1.37 |
| | PC2 | 3.15 | 1 | 0.0757 | -0.99 | -2.35 | 0.10 |
| | PC3 | 0.27 | 1 | 0.6036 | 0.21 | -0.58 | 1.12 |
| | PC4 | 5.61 | 1 | 0.0178 | -1.19 | -2.42 | -0.19 |
| | PC5 | 0.75 | 1 | 0.3878 | 0.36 | -0.45 | 1.24 |
| | Exact temperature | 6.42 | 1 | 0.0113 | 1.76 | 0.36 | 3.61 |
| Hot & Wet | PC1 | 13.33 | 1 | 0.0003 | 2.03 | 0.77 | 3.95 |
| | PC2 | 0.40 | 1 | 0.5262 | -0.29 | -1.26 | 0.62 |
| | PC3 | 6.14 | 1 | 0.0132 | -1.01 | -1.98 | -0.20 |
| | PC4 | 3.76 | 1 | 0.0524 | -0.87 | -1.94 | 0.01 |
| | PC5 | 2.03 | 1 | 0.1543 | -0.64 | -1.63 | 0.24 |
| | Exact temperature | 5.30 | 1 | 0.0214 | 1.48 | 0.19 | 3.33 |

**Table S4.** Factor loadings for the phylogenetic principal component analysis. Bioclimatic variables were extracted at the geographic coordinates for all taxa represented in our phylogenetic comparative analyses. The proportion of variance explained by each factor is given in parentheses.

| Bioclimatic variable | pPC1 (37.0%) | pPC2 (35.1%) |
|---|---|---|
| Mean temperature of the wettest quarter | 0.85 | -0.06 |
| Maximum temperature of the warmest month | 0.80 | -0.24 |
| $Log_e$(Precipitation of the wettest quarter) | 0.34 | 0.79 |
| $Log_e$(Precipitation of the driest month) | -0.04 | 0.85 |

**Table S5.** Effects of climatic temperature (pPC1), precipitation (pPC2), and their interaction on the likelihood that *Schizocosa* lineages evolve both complex courtship behaviors and exaggerated ornaments. Results were obtained from a phylogenetic logistic regression. The model estimated $\alpha = 3.7$ (rate of adaptation in Ornstein-Uhlenbeck model). Significance was determined by testing if the confidence intervals from 1000 bootstrapped parameter estimates overlapped zero.

| Response | Variable | $\beta \pm$ SE | Lower CI 95% | Upper CI 95% |
|---|---|---|---|---|
| Total display complexity $Log_e(+1)$ | pPC1 | 1.21 ± 0.57 | 0.26 | 2.88 |
| | pPC2 | 0.71 ± 0.39 | 0.01 | 1.86 |
| | pPC1 x pPC2 | 0.22 ± 0.56 | -1.17 | 1.42 |

**Table S6.** Model comparisons to determine how to incorporate phylogenetic signal in the phylogenetic generalized least squares analyses (PGLS). Models fit with Pagel's $\lambda$ did not converge. Preferred models are indicated in bold.

| Model formula | Rank | Model of evolution | df | AICc | ΔAICc | Weight |
|---|---|---|---|---|---|---|
| $Log_e$(Total display complexity + 1) ~ pPC1 + pPC2 + pPC1*pPC2 | **1** | **No phylogeny** | **5** | **104.92** | **0** | **0.93** |
| | 2 | Brownian motion | 5 | **110.02** | **5.10** | 0.07 |

**Table S7.** Effects of climatic temperature (pPC1), precipitation (pPC2), and their interaction on the evolution of total display complexity. Results were obtained from the best fitting phylogenetic generalized least squares models (PGLS). Significance was assessed using likelihood ratio tests comparing a full model to a model in which the variable of interest was removed. Standardized parameter estimates ($\beta$) and confidence intervals were obtained from the full models.

| Response | Variable | LR $\chi_1^2$ | df | P | $\beta$ | CI 95% |
|---|---|---|---|---|---|---|
| Total display complexity $Log_e(+1)$ | pPC1 | 4.83 | 1 | 0.0279 | 0.23 | 0.02 to 0.44 |
| | pPC2 | 14.64 | 1 | <0.0001 | 0.43 | 0.22 to 0.65 |
| | pPC1 x pPC2 | 0.13 | 1 | 0.7136 | 0.04 | -0.20 to 0.28 |

**Table S8.** Results from phylogenetic path analysis. Models are hypothesized evolutionary relationships among the number of distinct courtship behaviors, standardized ornament exaggeration score, pPC1 (temperature), and pPC2 (precipitation). Models assume evolution under Brownian motion. Preferred models are indicated in bold. Model type indicates whether trait exaggeration was binary (determined to be "complex" traits if the taxon expressed at least the median trait value for each trait; trait shading in Figure 4A) or continuous (trait bars in Figure 4A)

| Model type | Model | C | p | CICc | ΔCICc | Weight |
|---|---|---|---|---|---|---|
| Binary behavioral complexity and ornament exaggeration | (Figure S8C) Climate effects on ornament exaggeration emerge through behavior-ornament coevolution. | **2.27** | **0.8935** | **19.56** | **0.00** | **0.80** |
| | (Figure S8E) Climate effects on ornament exaggeration emerge through a combination of independent effects and behavior-ornament coevolution. | 0.21 | 0.9008 | 23.83 | 4.27 | 0.09 |
| | (Figure S8F) Climate effects on behavioral complexity emerge through a combination of independent effects and behavior-ornament coevolution. | 0.21 | 0.9008 | 23.83 | 4.27 | 0.09 |
| | (Figure S8B) Climate effects on behavioral complexity are independent of trait coevolution. | 10.07 | 0.1218 | 27.36 | 7.80 | 0.02 |
| | (Figure S8D) Climate effects on behavioral complexity emerge through behavior-ornament coevolution. | 16.00 | 0.0138 | 33.29 | 13.73 | 0.00 |
| | (Figure S8A) Climate effects on ornament exaggeration are independent of trait coevolution. | 23.31 | 0.0007 | 40.61 | 21.05 | 0.00 |
| Continuous behavioral complexity and ornament exaggeration | (Figure S9A) Climate effects on ornament exaggeration are independent of trait coevolution. | **8.72** | **0.1897** | **26.02** | **0.00** | **0.28** |
| | (Figure S9E) Climate effects on ornament exaggeration emerge through a combination of independent effects and behavior-ornament coevolution. | **2.53** | **0.2817** | **26.16** | **0.14** | **0.26** |
| | (Figure S9F) Climate effects on behavioral complexity emerge through a combination of independent effects and behavior-ornament coevolution. | **2.53** | **0.2817** | **26.16** | **0.14** | **0.26** |
| | (Figure S9B) Climate effects on behavioral complexity are independent of trait coevolution. | **9.37** | **0.1536** | **26.67** | **0.65** | **0.20** |
| | (Figure S8D) Climate effects on behavioral complexity emerge through behavior-ornament coevolution. | 16.99 | 0.0093 | 34.28 | 8.26 | 0.00 |
| | (Figure S8C) Climate effects on ornament exaggeration emerge through behavior-ornament coevolution. | 17.78 | 0.0068 | 35.08 | 9.06 | 0.00 |

**Table S9.** Varimax-rotated factor loadings for the principal component analysis of bioclimatic variables among *Schizocosa* assemblages. Bioclimatic variables for each assemblage were averaged over the entire assemblage area. The proportion of variance explained by each factor before rotation is given in parentheses (values for sliding window and non-overlapping grid cells on the left and right, respectively).

| Grid type | Bioclimatic variable | PC1 (51%; 50%) | PC2 (32%; 34%) |
|---|---|---|---|
| Sliding window | Mean temperature of the wettest quarter | 0.08 | 0.87 |
| | Maximum temperature of the warmest month | 0.12 | 0.88 |
| | $Log_e$(Precipitation of the wettest quarter) | 0.94 | -0.04 |
| | $Log_e$(Precipitation of the driest month) | 0.89 | 0.27 |
| Non-overlapping | Mean temperature of the wettest quarter | 0.08 | 0.88 |
| | Maximum temperature of the warmest month | 0.08 | 0.90 |
| | $Log_e$(Precipitation of the wettest quarter) | 0.94 | -0.06 |
| | $Log_e$(Precipitation of the driest month) | 0.89 | 0.28 |

**Table S10.** Model comparisons to determine how to incorporate spatial correlation structure in our assemblage-level spatially explicit generalized least squares analyses. Preferred models are indicated in bold.

| Grid type | Rank | Spatial correlation structure | df | AICc | ΔAICc | Weight |
|---|---|---|---|---|---|---|
| Sliding window | **1** | **Rational quadratic** | **8** | **953.52** | **0.00** | **0.91** |
| | 2 | Exponential | 8 | 958.16 | 4.64 | 0.09 |
| | 3 | Spherical | 8 | 965.87 | 12.34 | 0.00 |
| | 4 | Linear | 8 | 985.07 | 31.55 | 0.00 |
| | 5 | Gaussian | 8 | 987.59 | 34.06 | 0.00 |
| | 6 | No correlation | 7 | 1195.75 | 242.23 | 0.00 |
| Non-overlapping | **1** | **Rational quadratic** | **8** | **337.65** | **0.00** | **0.26** |
| | 2 | Exponential | 8 | 337.70 | 0.05 | 0.26 |
| | 3 | Gaussian | 8 | 338.92 | 1.27 | 0.14 |
| | 4 | Linear | 8 | 338.99 | 1.34 | 0.13 |
| | 5 | Spherical | 8 | 338.99 | 1.34 | 0.13 |
| | 6 | No correlation | 7 | 340.24 | 2.58 | 0.07 |

**Table S11.** Effects of an area's climatic temperature (PC2), precipitation (PC1), species richness, and sampling effort on the mean total display complexity score among species in the local assemblage. Results were obtained from the best fitting spatially explicit generalized least squares models. Significance was assessed using likelihood ratio tests comparing a full model to a model in which the variable of interest was removed. Standardized parameter estimates ($\beta$) and confidence intervals were obtained from the full models. Null $\beta$ upper 95% quantile levels were obtained from null distributions of 100 randomly simulated parameter estimates.

| Grid type | Variable | LR $\chi^2_1$ | df | P | $\beta$ | ±SE | Lower CL 2.5% | Upper CL 97.5% | Null $\beta$ 95% quantile |
|---|---|---|---|---|---|---|---|---|---|
| Sliding window | PC2 | 22.39 | 1 | <0.0001 | 0.32 | 0.06 | 0.19 | 0.44 | 0.38 |
| | PC1 | 35.65 | 1 | <0.0001 | 0.51 | 0.08 | 0.36 | 0.66 | 0.50 |
| | PC2xPC1 | 8.10 | 1 | 0.0044 | 0.15 | 0.05 | 0.05 | 0.25 | 0.27 |
| | $Log_e$(Sampling effort) | 2.22 | 1 | 0.1365 | 0.06 | 0.04 | -0.02 | 0.14 | 0.13 |
| | $Log_e$(Species richness) | 1.28 | 1 | 0.2578 | 0.04 | 0.04 | -0.04 | 0.12 | 0.52 |
| Non-overlapping | PC2 | 10.83 | 1 | 0.0010 | 0.26 | 0.07 | 0.12 | 0.41 | 0.43 |
| | PC1 | 22.30 | 1 | <0.0001 | 0.59 | 0.08 | 0.43 | 0.75 | 0.49 |
| | PC2xPC1 | 5.66 | 1 | 0.0173 | 0.16 | 0.07 | 0.03 | 0.30 | 0.32 |
| | $Log_e$(Sampling effort) | 0.14 | 1 | 0.7044 | -0.03 | 0.07 | -0.18 | 0.11 | 0.14 |
| | $Log_e$(Species richness) | 1.85 | 1 | 0.1735 | 0.10 | 0.08 | -0.05 | 0.25 | 0.42 |

**Table S12.** Model comparisons to determine how to incorporate spatial and phylogenetic random effects in our analysis of species persistence in response to recent climate change. Models with spatial structure included a random effect of the grid cell in which persistence was measured. Models with phylogenetic structure included a random effect of species nested within the two major *Schizocosa* clades[34]. Model type indicates whether the data were limited to grid cells where temperature change since the 1970's-90's was between 0.5-1.0°C (75 grid cells), or whether all 96 grid cells were included. For models with ΔAICc < 2, the model with the fewest parameters was preferred (lowest degrees of freedom). Preferred models are indicated in bold.

| Model type | Fixed effects | Rank | Random effects | df | AICc | ΔAICc | Weight |
|---|---|---|---|---|---|---|---|
| Temp change between 0.5-1°C (75 cells; 12 species; 153 persistance measurements) | Temp change<br>Precip change<br>Complexity<br>Temp x Complex<br>Precip x Complex | **1** | **(Clade:Species)** | **7** | **205.25** | **0.00** | **0.56** |
| | | 2 | Grid ID + (Clade:Species) | 8 | 206.11 | 0.86 | 0.37 |
| | | 3 | No random effects | 6 | 210.05 | 4.80 | 0.05 |
| | | 4 | Grid ID | 7 | 211.61 | 6.36 | 0.02 |
| Temp change between -0.6–1.4°C (96 cells; 12 species; 189 persistance measurements) | Temp change<br>Precip change<br>Complexity<br>Temp x Complex<br>Precip x Complex | 2 | Grid ID + (Clade:Species) | 8 | 250.62 | 0.00 | 0.42 |
| | | **1** | **(Clade:Species)** | **7** | **250.90** | **0.28** | **0.37** |
| | | 3 | No random effects | 6 | 253.20 | 2.58 | 0.12 |
| | | 4 | Grid ID | 7 | 253.72 | 3.10 | 0.09 |

**Table S13.** Effects of behavioral courtship complexity and ornament exaggeration on whether species persisted in response to changes in temperature and precipitation since the 1970's-90's. Courtship complexity and ornament exaggeration in these models are two-level factor variables, with species scored as having high courtship complexity and ornament exaggeration if they had higher than the median trait values among the species used in our phylogenetic comparative analysis. Model type indicates whether the data were limited to grid cells where temperature change since the 1970's-90's was between 0.5-1.0°C (75 grid cells), or whether all 96 grid cells were included. Results are shown for the best fitting GLMM, and the full model with spatial (grid ID) and phylogenetic (species nested within clade) random effects. Significance was assessed using likelihood ratio tests comparing a full model to a model in which the variable of interest was removed.

| Model type | Random effects | Variable | LR $\chi^2_1$ | df | P | $\beta$ | ±SE | CI 95% |
|---|---|---|---|---|---|---|---|---|
| Temp change between 0.5-1°C (75 cells; 12 species; 153 persistance measures) | (Clade:Sp) **Preferred model** | Temp change | 2.58 | 1 | 0.108 | 1.26 | 0.43 | 0.42 to 2.1 |
| | | Precip change | 0.01 | 1 | 0.935 | 0.74 | 0.34 | 0.08 to 1.41 |
| | | Complexity | 0.84 | 1 | 0.360 | 0.70 | 0.69 | -0.65 to 2.06 |
| | | Temp x Complex | 6.96 | 1 | 0.008 | -1.18 | 0.48 | -2.12 to -0.24 |
| | | Precip x Complex | 6.31 | 1 | 0.012 | -1.09 | 0.45 | -1.98 to -0.2 |
| | Grid ID + (Clade:Sp) | Temp change | 2.29 | 1 | 0.130 | 1.37 | 0.49 | 0.41 to 2.32 |
| | | Precip change | 0.01 | 1 | 0.926 | 0.88 | 0.41 | 0.08 to 1.68 |
| | | Complexity | 0.77 | 1 | 0.379 | 0.77 | 0.78 | -0.76 to 2.3 |
| | | Temp x Complex | 6.63 | 1 | 0.010 | -1.27 | 0.53 | -2.31 to -0.22 |
| | | Precip x Complex | 7.19 | 1 | 0.007 | -1.31 | 0.55 | -2.39 to -0.23 |
| Temp change between -0.6–1.4°C (96 cells; 12 species; 189 persistance measures) | (Clade:Sp) **Preferred model** | Temp change | 4.03 | 1 | 0.045 | 10.05 | 3.44 | 3.29 to 16.80 |
| | | Precip change | 0.07 | 1 | 0.790 | 0.10 | 0.05 | 0.01 to 0.20 |
| | | Complexity | 0.49 | 1 | 0.485 | 8.19 | 2.91 | 2.48 to 13.89 |
| | | Temp x Complex | 8.42 | 1 | 0.004 | -9.21 | 3.52 | -16.10 to -2.31 |
| | | Precip x Complex | 7.56 | 1 | 0.006 | -0.16 | 0.06 | -0.27 to -0.04 |
| | Grid ID + (Clade:Sp) | Temp change | 3.27 | 1 | 0.070 | 3.72 | 1.34 | 1.09 to 6.36 |
| | | Precip change | 0.04 | 1 | 0.836 | 0.87 | 0.40 | 0.10 to 1.64 |
| | | Complexity | 0.35 | 1 | 0.553 | 1.59 | 0.83 | -0.04 to 3.22 |
| | | Temp x Complex | 7.90 | 1 | 0.005 | -3.43 | 1.36 | -6.09 to -0.76 |
| | | Precip x Complex | 8.53 | 1 | 0.003 | -1.29 | 0.49 | -2.26 to -0.33 |

**Table S14.** Estimated marginal effects of behavioral courtship complexity and ornament exaggeration on whether species persisted in response to changes in temperature and precipitation since the 1970's-90's. Species' display complexity in these models is a two-level factor variable, with species scored as having high display complexity if they expressed at least the median behavioral complexity and ornament exaggeration values among the species used in our phylogenetic comparative analysis. Model type indicates whether the data were limited to grid cells where temperature change since the 1970's-90's was between 0.5-1.0°C (75 grid cells), or whether all 96 grid cells were included. Results are shown for the best fitting GLMM.

| Model type | Variable | Temp. $\beta$ | Temp. ±SE | Temp. CI 95% | Precip. $\beta$ | Precip. ±SE | Precip. CI 95% |
|---|---|---|---|---|---|---|---|
| Temp change between 0.5-1°C (75 cells; 12 species; 153 persistance measurements) | Complex displays | 1.26 | 0.43 | 0.42 to 2.10 | 0.74 | 0.34 | 0.08 to 1.41 |
| | Simple displays | 0.08 | 0.22 | -0.35 to 0.51 | -0.35 | 0.29 | -0.92 to 0.22 |
| Temp change between -0.6–1.4°C (96 cells; 12 species; 189 persistance measurements) | Complex displays | 3.40 | 1.17 | 1.12 to 5.60 | 0.71 | 0.33 | 0.07 to 1.35 |
| | Simple displays | 0.28 | 0.21 | -0.12 to 0.69 | -0.36 | -0.36 | -0.83 to 0.11 |

**Table S15.** Trait descriptions for each taxon used in the phylogenetic comparative analyses. Ornament exaggeration scores were obtained by Starrett and colleagues[34]. Behavioral courtship complexity scores were derived by counting the number of distinct courtship behaviors in the courtship descriptions and videos published in peer-reviewed articles. Courtship behaviors were delineated by distinct production mechanisms, and a single behavior could produce only visual, only vibrational, or simultaneous visual and vibrational stimuli. Distinct behaviors are separated by semicolons. References for courtship behaviors include the name, year, and journal abbreviation of articles cited in the main text reference list.

| Taxon | Ornament score | Courtship complexity | Visual-only behaviors | Vibration-only behaviors | Simultaneous visual & vibration behaviors | Courtship complexity reference |
|---|---|---|---|---|---|---|
| *S. minnesotensis* CO (simple) | 4 | 2 | leg waving | stridulation | NA | Starrett et al 2021 ISD |
| *S. salsa* NC (complex) | 3 | 4 | leg waving; leg arching | tremulation; stridulation | NA | Starrett et al 2021 ISD |
| *S. avida* OH (simple) | 0 | 2 | leg taps while walking | stridulation | NA | Starrett et al 2021 ISD |
| *S. mccooki* UT (simple) | 0 | 2 | leg quivering | palp drumming | NA | Starrett et al 2021 ISD |
| *S. mccooki* North CA (simple) | 0 | 2 | leg quivering | palp drumming | NA | Starrett et al 2021 ISD |
| *S. mccooki* South CA 1 (simple) | 0 | 2 | leg quivering | palp drumming | NA | Starrett et al 2021 ISD |
| *S. mccooki* South CA 2 (simple) | 0 | 2 | leg quivering | palp drumming | NA | Starrett et al 2021 ISD |
| *S. mccooki* South CA 3 (simple) | 0 | 2 | leg quivering | palp drumming | NA | Starrett et al 2021 ISD |
| *S. retrorsa* NC (complex) | 2 | 4 | push up; leg waving | palp drumming | leg tap | Hebets et al. 2008 AB; Choi & Hebets 2021 JA; Starrett et al 2021 ISD |

| | | | | | | |
|---|---|---|---|---|---|---|
| *S. retrorsa* MS (complex) | 2 | 4 | push up; leg waving | palp drumming | leg tap | Hebets et al. 2008 AB; Choi & Hebets 2021 JA; Starrett et al 2021 ISD |
| *S. chiricahua* AZ (simple) | 0 | 2 | NA | palp drumming | multi-leg movements with sound production | Starrett et al 2021 ISD |
| *S. aulonia* IN 1 (complex) | 4 | 4 | leg extend | palp drumming | leg percussion | Starrett et al 2021 ISD |
| *S. mimula* CO 1 (simple) | 0 | 2 | NA | palp drumming | leg strike | Starrett et al 2021 ISD |
| *S. mimula* CO 2 (simple) | 0 | 2 | NA | palp drumming | leg strike | Starrett et al 2021 ISD |
| *S. communis* ONT (simple) | 1 | 2 | multi-leg movements | stridulation | NA | Starrett et al 2021 ISD |
| *S. mccooki* ID (simple) | 2 | 2 | multi-leg movements | palp drumming | NA | Stratton & Lowrie 1984 JA; Starrett et al 2021 ISD |
| *S. mccooki* CO (simple) | 0 | 2 | multi-leg movements | palp drumming | NA | Stratton & Lowrie 1984 JA; Starrett et al 2021 ISD |
| *S. mccooki* SD (simple) | 0 | 2 | leg quivering | stridulation | NA | Starrett et al 2021 ISD |
| *S. cespitum* MB (simple) | 0 | 2 | multi-leg movements | stridulation | NA | Starrett et al 2021 ISD |
| *S. cespitum* SK (simple) | 2 | 2 | multi-leg movements | stridulation | NA | Starrett et al 2021 ISD |
| *S. humilis* MS (simple) | 1 | 5 | leg quivering; leg flicks | rapid stridulation; pulse stridulation | alternating leg strikes | McGinley et al. 2023 AN |
| *S. humilis* GA (complex) | 2 | 5 | leg quivering; leg flicks | rapid stridulation; pulse stridulation | alternating leg strikes | McGinley et al. 2023 AN |
| *S. humilis* FL (complex) | 2 | 5 | leg quivering; leg flicks | rapid stridulation; pulse stridulation | alternating leg strikes | McGinley et al. 2023 AN |

| | | | | | | |
|---|---|---|---|---|---|---|
| *S. duplex*<br>MS<br>(complex) | 2 | 3 | pre-mount leg flex | rapid stridulation; long pulse stridulation | NA | McGinley et al. 2023 AN |
| *S. crassipalpata*<br>SD<br>(simple) | 0 | 2 | NA | palp stridulation | leg tapping | Hebets et al. 2021 EE |
| *S. bilineata*<br>OH<br>(simple) | 6 | 2 | NA | palp stridulation | leg tapping | Hebets et al. 2021 EE |
| *S. segregata*<br>FL<br>(complex) | 6 | 4 | leg waving | stridulation; palp drumming | leg flicks | McGinley et al. 2023 AN |
| *S. saltatrix*<br>MS<br>(simple) | 1 | 3 | leg arch | stridulation; tremulation | NA | Stratton 2005 JA; Hebets et al. 2013 BES |
| *S. saltatrix*<br>KY<br>(simple) | 1 | 3 | leg arch | stridulation; tremulation | NA | Stratton 2005 JA; Hebets et al. 2013 BES |
| *S. floridana*<br>FL 1<br>(complex) | 2 | 3 | NA | palp stridulation; tremulation | leg taps | Rundus et al. 2011 Evol; Rosenthal et al. 2018 Evol |
| *S. floridana*<br>FL 2<br>(complex) | 2 | 3 | NA | palp stridulation; tremulation | leg taps | Rundus et al. 2011 Evol; Rosenthal et al. 2018 Evol |
| *S. crassipes*<br>NC<br>(complex) | 6 | 5 | bounce; extension; arch; wave | stridulation | NA | Stratton 2005 JA |
| *S. crassipes*<br>MS<br>(complex) | 6 | 5 | bounce; extension; arch; wave | stridulation | NA | Miller et al. 1998 AB; Stratton 2005 JA |
| *S. uetzi*<br>SC<br>(complex) | 2 | 3 | leg arching | two stridulatory behaviors | NA | Stratton 2005 JA; Hebets et al. 2013 BES; Hebets et al. 2005 BE |
| *S. stridulans*<br>KY<br>(complex) | 5 | 4 | leg arch | three stridulatory behaviors | NA | Hebets et al. 2008 AB; Hebets et al. 2013 BES |
| *S. ocreata*<br>FL<br>(complex) | 7 | 5 | leg taps; leg arch | tremulation; stridulation | cheliceral strike | Stratton 2005 JA |

| | | | | | | |
|---|---|---|---|---|---|---|
| *S. rovneri*<br>IL<br>(simple) | 0 | 2 | bounce | palp stridulation during bounce | NA | Davis 1989 AMN |
| *S. rovneri*<br>MS<br>(simple) | 0 | 2 | bounce | palp stridulation during bounce | NA | Stratton & Uetz 1986 Evol |
| *S. ocreata*<br>MS<br>(complex) | 7 | 5 | leg taps; leg arch | tremulation; stridulation | cheliceral strike | Fowler-Finn et al. 2015 CZ |
| *S. ocreata*<br>CO<br>(complex) | 8 | 5 | leg taps; leg arch | tremulation; stridulation | cheliceral strike | This Study; McGinley et al. 2023 AN |
| *S. ocreata*<br>TN<br>(complex) | 9 | 5 | leg taps; leg arch | tremulation; stridulation | cheliceral strike | This Study |
| *S. ocreata*<br>WV<br>(complex) | 9 | 4 | leg taps; leg arch | stridulation | cheliceral strike | Gordon & Uetz 2011 AB; Stratton & Uetz 1986 Evol |

**Table S16.** The display traits and phenological periods used for each species in the analysis of species persistence. Ornament and courtship complexity scores are averages across all of the species' corresponding taxa in the phylogenetic comparative analyses. Display complexity categories are in parentheses, with species scored as having high display complexity if they expressed at least the median behavioral complexity and ornament exaggeration values among the species used in our phylogenetic comparative analysis. The month ranges represent the months of the year where each species is most reproductively active, determined by collection dates in peer-reviewed articles and/or verified by examining monthly iNaturalist observation frequencies. The month ranges were used to calculate change in temperature and precipitation since the 1970's-90's during the reproductive period for each species.

| Taxon | Ornament score | Courtship complexity | Clade | Month of reproductive activity | Reference for months reproductively active |
|---|---|---|---|---|---|
| *S. mccooki* (simple) | 0.25 | 2 | "mccooki" | May-July | Stratton & Lowrie 1984 JA |
| *S. retrorsa* (complex) | 2 | 4 | "mccooki" | March-May | Choi & Hebets 2021 JA |
| *S. avida* (simple) | 0 | 2 | "mccooki" | April-June | iNaturalist only |
| *S. minnesotensis* (simple) | 4 | 2 | "mccooki" | March-May | iNaturalist only |
| *S. ocreata* (complex) | 8 | 4.8 | "ocreata" | April-June | This Study; McGinley et al. 2023 AN |
| *S. rovneri* (simple) | 0 | 2 | "ocreata" | March-May | iNaturalist only |
| *S. crassipes* (complex) | 6 | 5 | "ocreata" | April-June | iNaturalist only |
| *S. saltatrix* (simple) | 1 | 3 | "ocreata" | March-May | iNaturalist only |
| *S. bilineata* (simple) | 6 | 2 | "ocreata" | April-June | Hebets et al. 2021 EE |
| *S. crassipalpata* (simple) | 0 | 2 | "ocreata" | March-May | Hebets et al. 2021 EE |
| *S. duplex* (complex) | 2 | 3 | "ocreata" | April-June | McGinley et al. 2023 AN |
| *S. uetzi* (complex) | 2 | 3 | "ocreata" | May-July | Hebets et al. 2005 BE |
